# Mechanical imbalance between normal and transformed cells drives epithelial homeostasis through cell competition

**DOI:** 10.1101/2023.09.27.559723

**Authors:** Praver Gupta, Sayantani Kayal, Nobuyuki Tanimura, Shilpa P. Pothapragada, Harish K. Senapati, Padmashree Devendran, Yasuyuki Fujita, Dapeng Bi, Tamal Das

## Abstract

Cell competition in epithelial tissue eliminates transformed cells expressing activated oncoproteins to maintain epithelial homeostasis. Although the process is now understood to be of mechanochemical origin, direct mechanical characterization and associated biochemical underpinnings are lacking. Here, we employ tissue-scale stress and compressibility measurements and theoretical modeling to unveil a mechanical imbalance between normal and transformed cells, which drives cell competition. In the mouse intestinal epithelium and epithelial monolayer, transformed cells get compacted during competition. Stress microscopy reveals an emergent compressive stress at the transformed loci leading to this compaction. A cell-based self-propelled Voronoi model predicts that this compressive stress originates from a difference in the collective compressibility of the competing populations. A new collective compressibility measurement technique named gel compression microscopy then elucidates a two-fold higher compressibility of the transformed population than the normal population. Mechanistically, weakened cell-cell adhesions due to reduced junctional abundance of E-cadherin in the transformed cells render them collectively more compressible than normal cells. Taken together, our findings unveil a mechanical basis for epithelial homeostasis against oncogenic transformations with implications in epithelial defense against cancer.

## Introduction

Epithelial homeostasis is crucial for maintaining tissue integrity and function, and it is achieved through a balance of cell proliferation, differentiation, and extrusion [1–4]. Epithelial cells actively extrude unfit cells that may compromise the epithelial barrier and tissue integrity. For example, cells under metabolic stress or exhibiting DNA damage are identified and expelled by their neighbors, preserving overall tissue health [1]. Additionally, cells with disrupted polarity or adhesion properties are removed via a similar extrusion process [5]. This selective extrusion is regulated by signals from surrounding healthy cells, ensuring the elimination of potentially disruptive cells and maintaining epithelial homeostasis. Furthermore, the process is not only limited to cells with apparent damage but also includes cells that are simply older or less fit compared to their neighbors [1, 6]. These mechanisms collectively ensure that the epithelial layer remains robust and functional, capable of serving as a barrier and interface between the external environment and the body’s internal milieu. Through these processes, epithelial tissues also protect themselves from potential damage, prevent the onset of disease, and ensure long-term functionality and resilience.

Importantly, there is a broad class of non-cell autonomous cell-elimination processes called cell competition, which maintains epithelial homeostasis during organ development and ageing [7, 8]. In addition, cell competition mediates a non-cell autonomous removal of transformed mutant cells expressing constitutively active forms of oncoproteins such as HRas^V12^ [9–12], thus maintaining epithelial tissue homeostasis and integrity against oncogenic transformations. This process is often referred to as epithelial defense against cancer (EDAC). In this process, the transformed cells that are surrounded by normal epithelial cells, get eliminated from the tissue by apical extrusion or by basal delamination [11]. Despite some recent progress, the mechanisms underlying how normal cells recognize and eliminate the transformed cells remain largely elusive [9]. The competitive removal of transformed cells requires extensive reorganization of force-bearing cytoskeletal elements in the surrounding normal cells, particularly at the interface between normal and transformed cells [13–15]. The process additionally depends on the mechanics of the extracellular matrix [16], collectively indicating possible modulations of cellular mechanical forces [15, 17–20].

Mechanical forces play critical roles during cell competition in general [15, 17]. For example, a differential in cell proliferation potential between two competing populations can drive out the population with a lower proliferation potential by compaction. Also, cells carrying distinct genetic lesions can have varying sensitivities to compaction, which might lead to their elimination. These observations explain how a fast-growing population bearing higher resistance to deformation [15] and elimination can have a competitive advantage over a slow-growing population [18, 19, 21, 22]. However, it is not clear how cell elimination occurs when the proliferative differential is trivial. For example, in the competitive elimination of epithelial cells expressing oncoprotein HRas^V12^ by normal wildtype epithelial cells [11, 12, 16], the proliferation differential might not be a critical factor. Here, we probed the growth potential of these populations by quantitating their number expansion in time, starting from a similar number density (Supplementary Figs. 1A-B). We also directly measured the proliferation rate using ethynyl deoxyuridine (EdU) labeling (Supplementary Figs. 1C-D). These experiments revealed that the difference in growth potential between transformed and wildtype populations is not significant. If anything, HRas^V12^-expressing transformed cells show slightly higher proliferation post-48 hours of growth. Thus, in this model for cell competition, despite having a slightly higher growth rate, the transformed cells get eliminated in the presence of normal cells. The mechanisms underlying this competitive elimination of the transformed cells by normal cells remain poorly understood. While tissue and cell mechanics might play a crucial role here, in the absence of direct measurement of forces during cell competition, it is unclear how tissue mechanics regulate non-proliferative cell competition between normal and transformed cells in epithelia.

To understand how epithelial tissue eliminates HRas^V12^-expressing transformed cells, two inducible models have been pivotal – an *in vivo* mouse intestinal epithelium model [12] and an *in vitro* Madin-Darby canine kidney (MDCK) cell culture model [11]. The mouse model [12] involves a recombinant mouse strain viz. villin-HRas^V12^-GFP (villin-*Cre-ER^T2^*; *LSL-HRas^V12^-IRES-EGFP*), wherein intraperitoneal injection of tamoxifen leads to Cre recombinase-mediated induction of HRas^V12^ genetic recombination in select cells which then constitutively express HRas^V12^ in the intestinal epithelium (Fig. 1A). At the same time these transformed cells express EGFP and hence, can be tracked. In the cell culture model [11], wildtype and HRas^V12^-expressing MDCK epithelial cells are co-cultured (Fig. 1B and Supplementary Fig. 2A) leading up to a mosaic occurrence of transformed colonies surrounded by wildtype cells and thus, mimic competition. In this case, HRas^V12^ is tagged with GFP to mark the transformed cells. In this model, HRas^V12^ expression is dependent on a Tet-on system and induced by adding a tetracycline derivative, doxycycline to the culture medium. Here, we co-cultured wildtype and HRas^V12^-expressing transformed cells in a 40:1 ratio (Fig. 1B and Supplementary Fig. 2A) so that in the absence of doxycycline, as expected, transformed cells would form small colonies (2-10 cells), surrounded by wildtype cells. We then exposed this heterogeneous epithelial monolayer to a medium containing doxycycline, which induced HRas^V12^ expression in the transformed cells (Fig. 1B). Additionally, we employed 4 kPa polyacrylamide gels as the substrate for this model to mimic physiological stiffness [16]. While the mouse model of cell competition is more physiologically relevant than the cell culture model, the latter is more controllable than the former and amenable to mechanical measurements. In both models, HRas^V12^-expressing transformed cells were apically extruded out of the monolayer only when they were in contact with the surrounding normal cells (Figs. 1C-D), as has been reported previously [11], implying that the extrusion of HRas^V12^-expressing cells from heterogeneous epithelial tissue is a non-cell autonomous process driven by cell competition. From here onwards, we sort out to characterize the pre-extrusion mechano-phenotypes associated with cell competition during epithelial defense against cancer.

**Figure 1.**
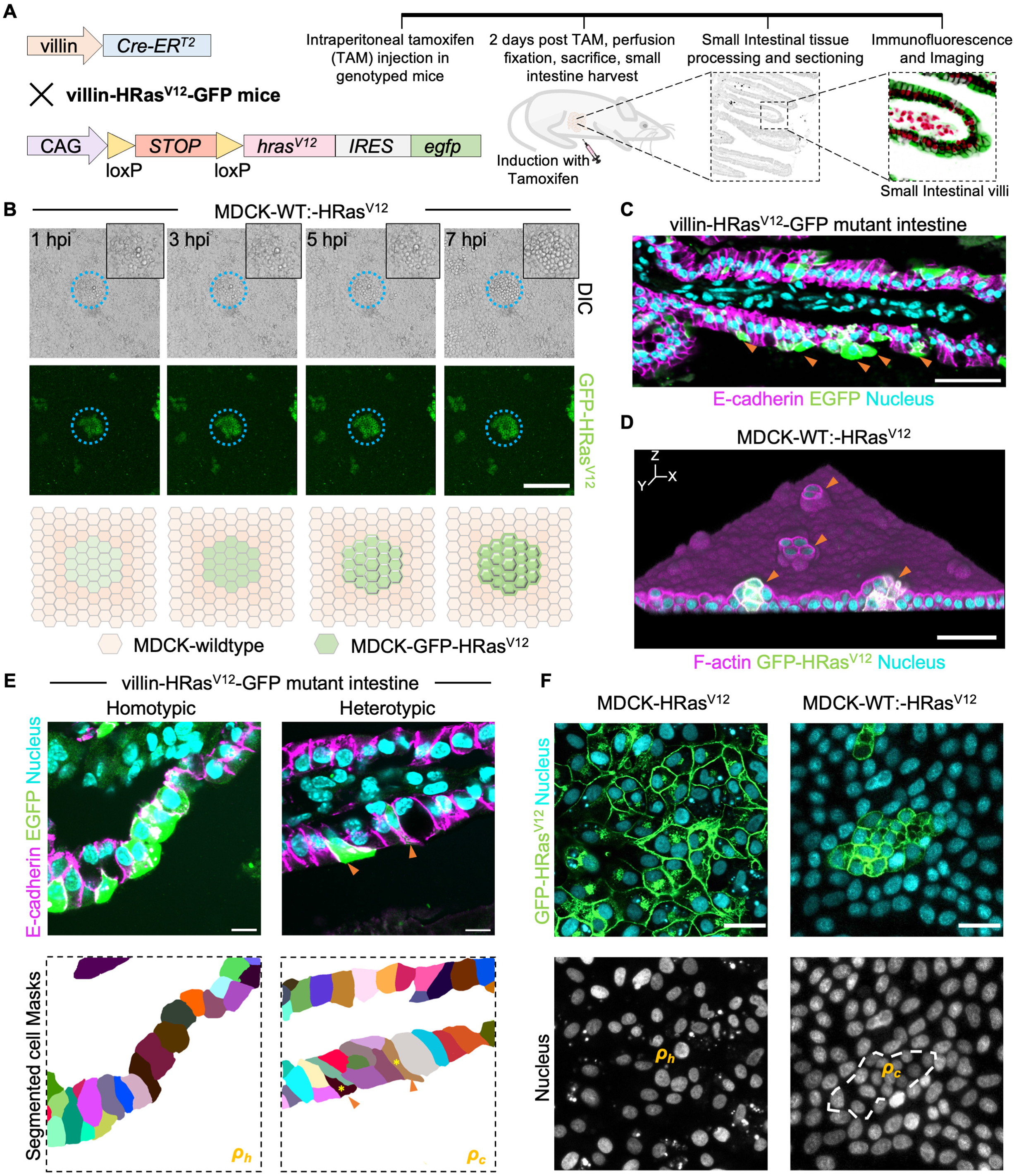
Competition induced compaction of HRas^V12^-transformed mutants *in vivo* and *in vitro*. **A.** Schematic representation of the genetic makeup and experimental workflow pertaining to the cell competition mice model. **B.** Heterogenous competing population constituted by wildtype and HRas^V12^-expressing transformed MDCK cells (40:1, number ratio) cultured on a 4 kPa polyacrylamide gel. Induction with doxycycline (dox) shows progressive GFP-HRas^V12^ expression and concomitantly, the transformed cells round up to finally extrude out of the monolayer (transformed colony is marked by blue dotted circle, also shown in the insets), hpi-hours post induction. Scale bar, 50µm. **C.** Representative immunofluorescence images of intestinal villi epithelium from villin-*Cre-ER^T^*^2^*; LSL*-*hras^V12^*-*IRES*-*egfp* (villin-HRas^V12^-GFP) mice after tamoxifen injection, as detailed in **A**, depicting selective competitive extrusion of HRas^V12^-expressing transformed (villin-HRas^V12^-GFP) but not wildtype cells. Scale bar, 50 µm. **D.** 3-dimensional rendered view of a heterogenous competing population of MDCK cells cultured over 4 kPa polyacrylamide gel. Orange arrow heads point towards the extruding transformed cell groups. Scale bar, 50 µm. **E.** cell spreading area preferred by HRas^V12^ and simultaneously EGFP expressing transformed cells in villin-HRas^V12^-GFP genetic background under homeostatic conditions (referred to as 𝛥_ℎ_) or under wildtype epithelium and transformed competition condition (referred to as 𝛥_𝑐_). Bottom panels depict the corresponding segmented cell masks. Scale bar, 10 µm. **F.** Cell spreading area preferred by HRas^V12^-transformed MDCK cells under homeostatic conditions (referred to as 𝛥_ℎ_) or under competition condition (referred to as 𝛥_𝑐_). Scale bar, 20 µm.

## Results

### Non-cell autonomous, competition-driven compaction of HRas^V12^-expressing cells

We initially observed that, in both models, the yet-to-be-extruded transformed cells appeared compacted when they were in competition with normal cells (Figs. 1E-F). First, in the intestinal epithelium of the mouse model, we compared the thickness of the HRas^V12^-expressing cells in the area where all cells expressed HRas^V12^ against the thickness of HRas^V12^-expressing cells in the area where these cells are sparsely distributed amongst normal cells (Fig. 1E). This comparison revealed that the thickness of HRas^V12^-expressing cells was reduced by more than 1.7-fold when they were surrounded by wildtype cells. These observations pointed towards a selective, competition-dependent compaction of HRas^V12^-expressing transformed cells but not control cells, in the intestinal villi of mice. Next, in the cell culture model, we compared the cell area of the HRas^V12^-expressing cells in homogeneous cultures versus in co-cultured systems where these cells were surrounded by wildtype cells. We found that in the homogeneous culture, surrounded by their own kind, HRas^V12^-expressing cells exhibited higher cell areas than even wildtype cells in a homogeneous normal epithelium. However, in the co-cultured system, in the presence of wildtype cells, the area of the HRas^V12^-expressing cells was reduced by more than 2.5 folds, as compared to their area under homogeneous conditions (Fig. 1F). Together these results established the generality of a competition-associated non-cell autonomous compaction of HRas^V12^-expressing cells, which might be driven by mechanobiological factors.

### Competition-induced compressive stress at the transformed mutant locus

Although, the mechanobiology of competition between wildtype and transformed cells remains elusive, there are indications that the core mechanoresponsive elements influencing cell and tissue mechanics could be involved in the competition-mediated extrusion of transformed cells [15, 16]. For example, at the locus where transformed cells interfaced wildtype cells in their vicinity, an active remodeling of the actomyosin occurred towards the transformed-wildtype interface, as highlighted by distinct phosphorylated myosin II light chain (pMLC2, active form of myosin II motor [23]) surrounding transformed cells (Supplementary Fig. 2B). However, how cellular forces change during cell competition remains unknown. We, therefore, elucidated the temporal evolution of mechanical force field in the cell culture model, by combining traction force microscopy with Bayesian inverse stress microscopy [24] (Figs. 2A-C). We also used Bayesian force inference [25] (Figs. 2D-F) in another independent set of experiments. From traction force microscopy experiments, we observed that upon induction of HRas^V12^-expression, traction forces increased along the interface between wildtype and transformed populations (Fig. 2A and Supplementary Fig. 2E). Simultaneously, stress microscopy revealed local compressive stress appearing in the regions occupied by transformed cells (Fig. 2C and Supplementary Fig. 2E). We concluded that this compressive stress is a unique mechanical signature of cell competition since in a homogeneous population of either normal or transformed cells, stresses remained predominantly tensile (Supplementary Figs. 2C-D), as reported previously [24, 26, 27]. While the complete extrusion of transformed cells took at least six hours after doxycycline induction (Fig. 1B and Supplementary Fig. 2E), compressive stresses appeared around two hours post-doxycycline exposure (Supplementary Fig. 2E), indicating that compressive stress precedes cell rounding up and extrusion (Supplementary video 1). In addition, compressive stress appeared to be restricted only to transformed cells as the background tissue comprising normal cells presented a predominantly tensile stress field (Fig. 2C, Supplementary Figs. 2C-D). To capture the cellular-level effect of the coarse-grained compressive stress obtained from the stress microscopy, we then used Bayesian force inference to compute relative interfacial tension at the cellular scale and relative intracellular pressure (Figs. 2E-G). This analysis revealed lower interfacial tension (Fig. 2E) and higher intracellular pressure (Figs. 2F-G) in transformed cells as compared to surrounding normal cells at the time when multicellular compressive stress appeared. Together these experiments revealed that while epithelial tissues are generally tensile systems, during the competition between normal and transformed cells, compressive stress arises. This compressive stress precedes the extrusion of transformed mutant cells and is a unique mechanical signature of cell competition.

**Figure 2.**
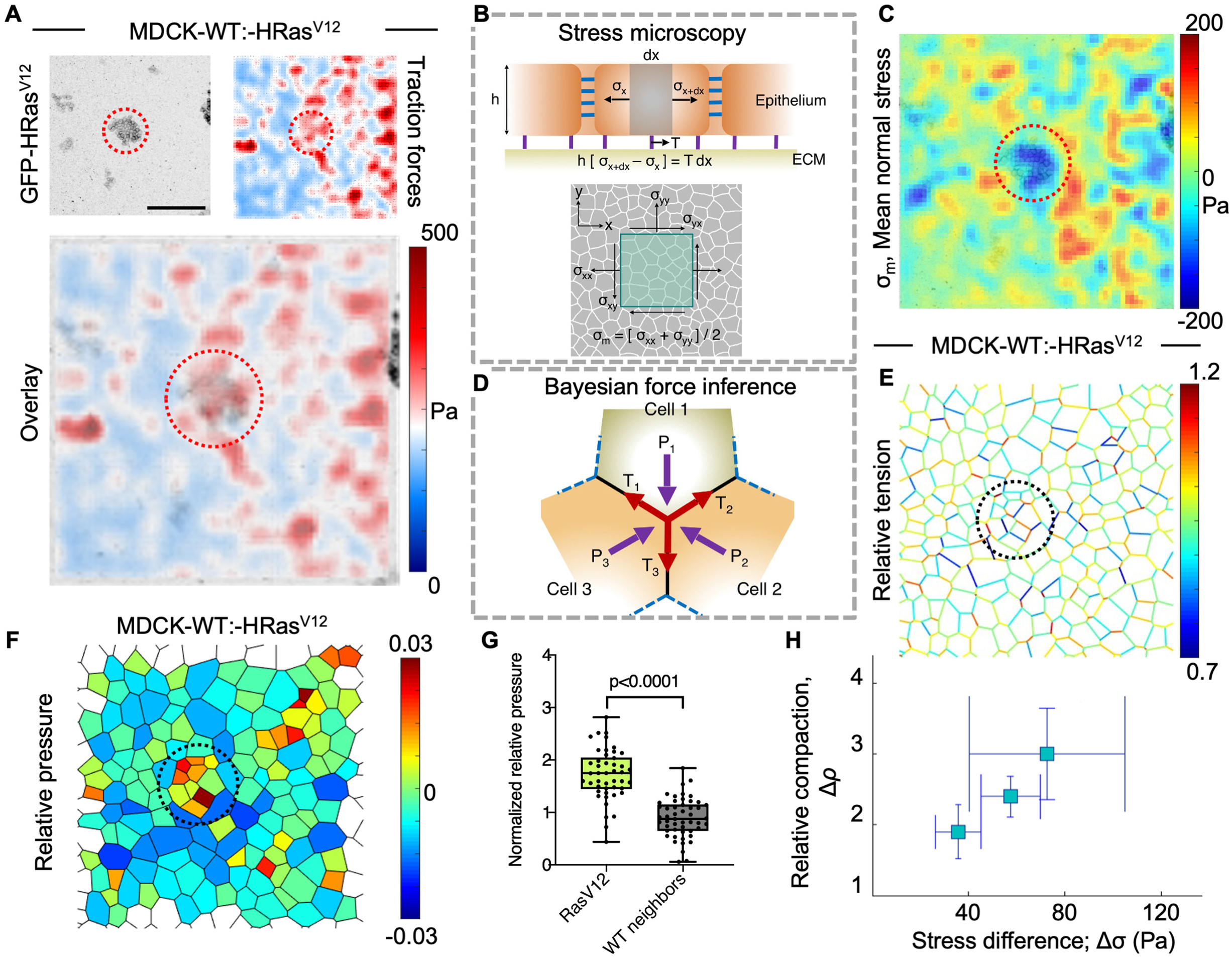
Competition dependent compressive stress at the HRas^V12^-transformed mutant locus. **A.** Traction force microscopy generated map of the forces exerted by MDCK-wildtype (-WT) cells during competition between HRas^V12^-transformed mutant MDCK and wildtype cells (transformed colony is marked by red dotted circle), analysis done post 2 hours of doxycycline (dox) induction. Scale bar, 50 µm. **B.** Infographic illustration describing the principle underlying monolayer stress microscopy. **C.** Stress microscopy generated mean normal stress map depicting compressive stress at the transformed loci (marked by red dotted circle) during competition between WT and MT cells, analysis done post 2 hours of induction with dox. Scale same as in **A**. **D.** Infographic illustration describing Bayesian force inference. Bayesian force inference generated **E.** interfacial tension map and **F.** relative intracellular pressure map during competition between WT and MT cells (transformed colony marked by black dotted circle), inference performed post 2 hours induction with dox. Scale same as in **A**. **G.** normalized relative intracellular pressure characterized with Bayesian force inference, plotted from a pool of 3 independent experiments, analysis performed post 2 hours induction with dox. Data are mean±sem. Statistical significance was calculated using Unpaired t-test with Welch’s correction. **H.** Relative compaction, 𝛥, defined as 𝛥 = (𝛥_𝑐_ – 𝛥_ℎ_)/ 𝛥_ℎ_ is determined for competing HRas^V12^-transformed cells under different times of induction (and thus experiencing different levels of compressive stress (plotted in Δσ). Data is represented as the relative compaction with the maximum compressive stress obtained for transformed colonies revealing a nearly linear relationship between compressive stress and relative compaction. Data are mean±sem from three independent experiments. Statistical significance was calculated using Unpaired t-test with Welch’s correction.

We estimated the competition-associated compaction relative to the homeostatic density by calculating relative compaction as Δρ = (*ρ_c_ − ρ_h_)/ρ_h_* (Fig. 2H, Eq. SI-6). We then hypothesized that this compaction could be a consequence of compressive stress and hence, compaction should increase with the increasing compressive stress. To test this hypothesis, we varied the concentration of doxycycline, which led to different levels of HRas^V12^-expression. Consequently, the compressive stress also changed, increasing with increasing doxycycline concentration (Supplementary Fig. 2E). Next, plotting the relative compaction with the maximum compressive stress obtained for HRas^V12^-expressing colonies revealed a nearly linear relationship between the compressive stress and the relative compaction (Fig. 2H), indicating a critical consequence of compressive stress on competition-associated compaction of the transformed cells.

### Destabilized adherens junctions in transformed mutant cells

We had previously elucidated one of the mechano-phenotypes pertaining to epithelial cell competition, which is the homeostatic preferred area (inverse of homeostatic density, the peak density of a confluent monolayer at which the cell division rate becomes negligible). To study the general effect of transformation on epithelial mechanics, we next resorted to adherens junctions in transformed cells both *in vivo* and *in vitro*, since the integrity of cell-cell junctions is key to epithelial function and physiology [28].

We henceforth looked at the integrity of cell-cell junctions in normal and transformed mutant cells. When we probed for endogenous E-cadherin in the *in vivo* mice intestinal epithelium harboring ubiquitous HRas^V12^ mutation, we found marked disruption in the junctional patterning of E-cadherin only in these mutants and not the control epithelium, supporting the phenotype that upon *hras* transformation, the homotypic adhesions are aberrant and occurs in physiology (Figs. 3A-D). We further characterized homotypic cell-cell adhesions in MDCK wildtype and transformed populations by immunostaining for E-cadherin (Figs. 3E-H). We found that in the transformed cells, as seen in the mice intestinal epithelium, E-cadherin was aberrantly localized to cell-cell junctions in addition to the vesicular distribution within the cytoplasm (Figs. 3F, H). This cytoplasmic vesicular E-cadherin pool was not observed in the wildtype cells (Fig. 3E, G), indicating an instability in cell-cell junctions in the transformed cells. We further corroborated this observation by performing super-resolution spinning-disk confocal microscopy using optical photon reassignment (SoRa) [29] with E-cadherin (stained by an antibody), both *in vivo* in the mice model of cell competition (Figs. 3C, D) and *in vitro* in MDCK cultures (Figs. 3I,J), to show that junctional E-cadherin was significantly deteriorated in the transformed cells as compared to the junctions of wildtype cells (Figs. 3C-D,I-J). Alongside, a marker protein for stable cell-cell junctions, alpha-catenin, showed a distinct cytoplasmic distribution pattern owing to the reduction in mature adherens junctions, only in the transformed cells (Supplementary Figs. 3A-D). This result reaffirmed the fact that stable cell-cell junctions were compromised in the transformed cells.

**Figure 3.**
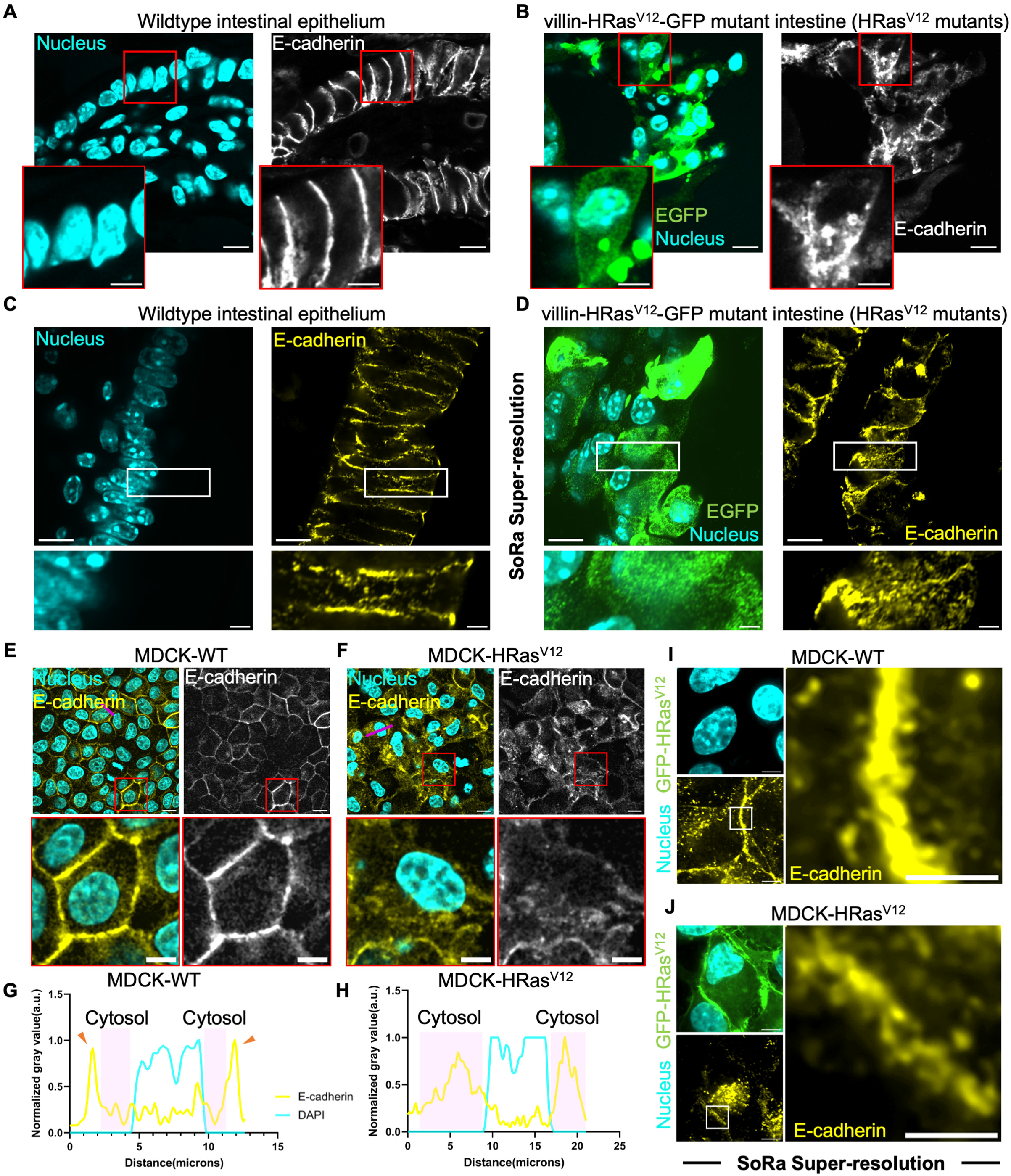
Perturbed adherens-based cell-cell junctions as distinct mechano-phenotype of HRas^V12^ transformed cells *in vivo* and *in vitro*. **A,B.** Representative immunofluorescence images of mice intestinal epithelium depicting endogenous E-cadherin in wildtype cells under villin-HRas^V12^-GFP background (**A**) or HRas^V12^ and simultaneously EGFP (*HRas^V12^-IRES-EGFP*) expressing cells in villin-HRas^V12^-GFP background (**B**). Insets are shown corresponding to each image from the two scenarios, Scale bar, 10 µm for all image panels and 2 µm for all insets. **C,D.** Super-resolution spinning-disk confocal microscopy using optical photon reassignment (SoRa) image panels to show endogenous E-cadherin in the intestinal villi epithelium in wildtype cells (**C**) or in Hras^V12^ and simultaneously EGFP expressing cells (**D**) under villin-HRas^V12^-GFP background. Insets are shown corresponding to each image from the two scenarios. Scale bar, 10 µm for all image panels and 2 µm for all insets. Representative panels to show immunostained endogenous E-cadherin in (**E**) MDCK-wildtype (-WT) and or (**F**) MDCK-HRas^V12^ transformed cells cultured in homogenous conditions. Insets are shown (marked by red boxes) to represent one cell in each condition. Scale bar, 10 µm for all image panels and 2 µm for all insets. **G,H.** Respective intensity line-plot (Magenta line refers to the intensity plot line) for WT (**G**) or transformed (**H**) cells. DAPI intensity (marked by cyan line) is plotted for reference to the nucleus. Pink boxes refer to the cytosol in each cell. Intensity values are normalized to the highest value within each dataset. E-cadherin shows distinct junctional accumulation in WT cells but diffused cytoplasmic signal in transformed cells. **I,J.** Super-resolution spinning-disk confocal microscopy using optical photon reassignment (SoRa) image panels to show endogenous E-cadherin in WT (**I**) or transformed (**J**) cells. Junctional E-cadherin is shown in insets from the two cell-types to depict E-cadherin loss in the transformed cells. Scale bar, 10 µm for all image panels and 2 µm for all insets.

Furthermore, live-cell imaging of junctional membranes in a monolayer of transformed cells displayed remarkably dynamic and continuous short-time-scale ruffling whilst junctional membranes in wildtype cells were stable at such short timescales and did not show this peculiar phenotype (Supplementary Figs. 3E-G). These results revealed instability in cell-cell junctions specifically in the transformed cells, while the normal cells showed relatively stable cell-cell adhesions. Surprisingly, we did not observe any significant difference in cell-matrix adhesions, as marked by endogenous paxillin in the two cell types (Supplementary Figs. 3H-J). Taken together, these results elucidate the characteristic differences between wildtype and transformed cell populations, which we subsequently feed into our computational model.

Having discovered fundamental mechanobiological differences that transformation incurs, we worked towards understanding why transformed cells experience compressive stress specifically during competition with the wildtype cells. To this end, we proposed a physical modeling of a tissue containing heterogeneous mixture of transformed and wildtype cells using the Self-Propelled Voronoi model [30], to accurately capture this heterogeneity in key differential attributes between wildtype and transformed populations.

### The computational Self-Propelled Voronoi model explains the origin of compressive stress

To gain a deeper understanding of the emergent mechanics resulting from competition, we used a well-known physical model for confluent tissue, the Self-Propelled Voronoi model (SPV) [28, 30–32], to simulate a heterogeneous monolayer. In this model, the heterogeneities in cell types were represented by differences in the single cell-level mechanical parameters that governed each cell (details of the model are outlined in the SI text). These parameters include the single-cell area and perimeter elastic moduli 𝐾_𝐴_ and 𝐾_𝑃_, and homeostatic preferred cell area and perimeter 𝐴_0_ and 𝑃_0_ respectively. We simulated a co-culture of two types of cells in the SPV, in which a circular island of transformed mutant cells (M) was surrounded by wildtype cells (WT) (Fig. 4A). In this study, we specifically investigated the emergent behaviour when these properties differed between transformed and wildtype cells, focusing on the differences in 𝐾_𝐴_, 𝐴_0_, and 𝑃_0_ values. In our simulations we therefore set 𝐾_𝑃_ = 1 for all cells and use *A*^*WT*^_0_ as the unit of length. Therefore, the sum total nondimensionalized mechanical energy function for the cell layer (derived from Eq. SI-1 in SI Section) is given by:

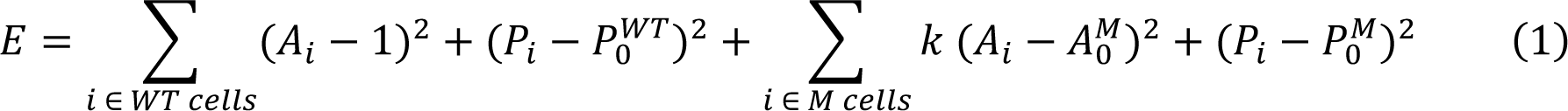

**Figure 4.**
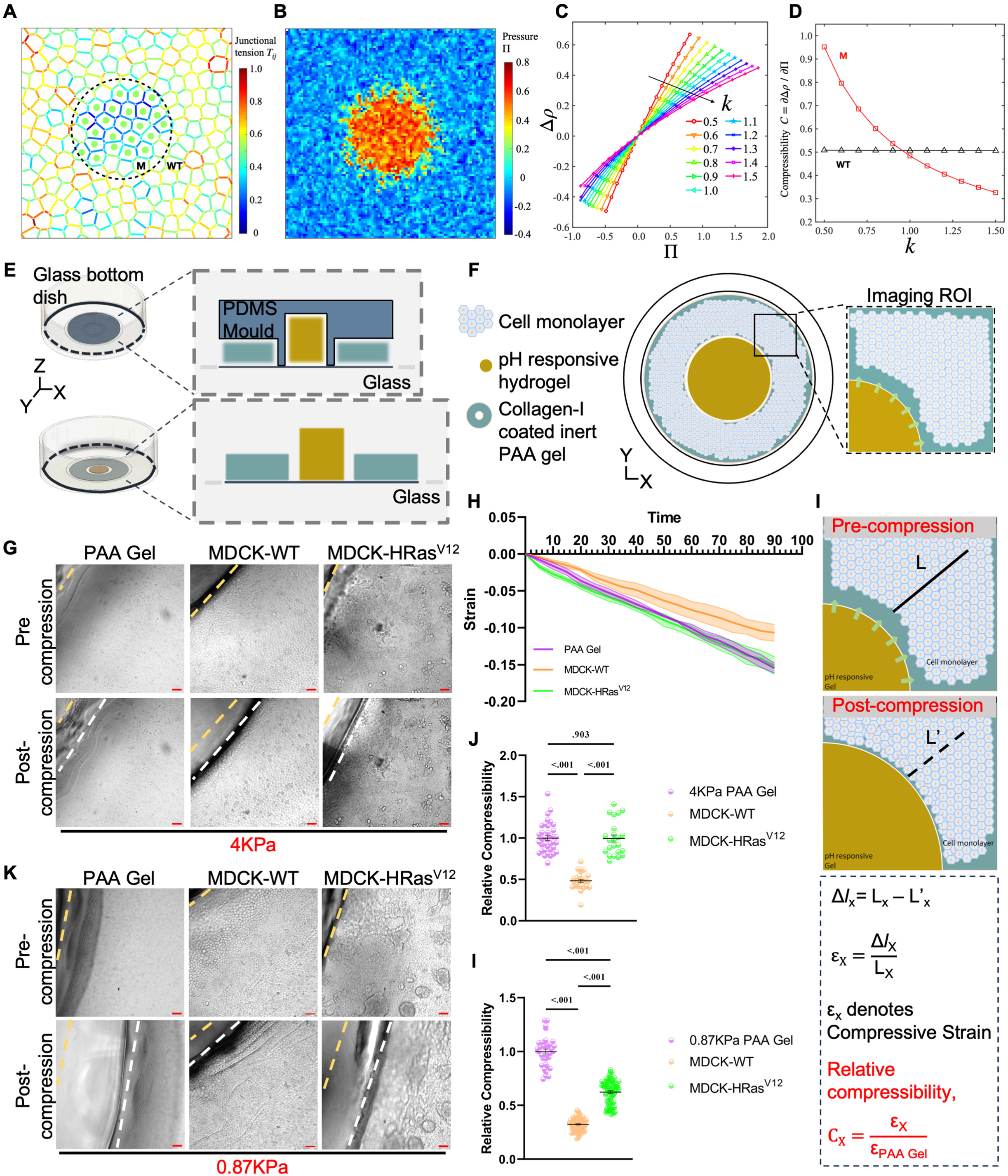
*In silico* self-propelled Voronoi model recapitulates competition-dependent mechanical features to discover differential collective compressibility, empirically tested by Gel Compression Microscopy. **A.** Junctional tension at the edges shared by cells in a circular island of transformed mutant (M) cells (which constitute 12% of the total number of cells in a randomly generated ensemble, with the island’s diameter being approximately 40% of the length of the box) is significantly lower than that of their surrounding wild-type (WT) counterparts. This is due to the single-cell parameters of the M cells, which include*K^M^_A_* = 0.7, *A^M^_0_* = 0.8, *P^M^_0_* = 3.8, which corresponds to a shape index of 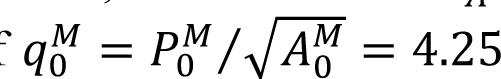 in contrast to the WT cells with parameters *K^WT^_A_* = 1.0, *A*^*WT*^_0_ = 1.0, *P^WT^_0_* = 3.65 (*i.e. q^WT^_0_* = 𝐴 0 0 0 3.65). As a result, the M colony exhibits lower edge tension and is less rigid than its surrounding WT cells. **B.** the average pressure in the M island (with the same spatial distribution as shown in Fig. 2F and coarse-grained over 50 ensembles of initial configurations) is elevated, with single-cell parameters of*K^M^_A_* = 0.7, *A^M^_0_* = 1.5, *P^M^_0_* = 3.8. In contrast, WT 𝐴 0 0 cells have the same single-cell parameters as in **A**. **C.** the relative compaction level, denoted by Δρ = *A*_0_/*Ā* − 1, of the M population is a function of the average pressure (Π) and depends on an array of cell area elasticity ratios, 𝑘 =*K^M^_A_*⁄*K^WT^_A_*. In the M island, an increase in the value 𝐴 𝐴 of Π is driven by an increase in *A^M^_0_*. **D.** the linear-response compressibility, denoted by 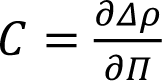, for M and WT cells as a function of 𝑘. **E,F.** Conceptual representation depicting the **E**, basis and functionality, and **F**, the data acquisition as done when performing Gel Compression Microscopy (GCM). Micropatterned PDMS mould (shown in blue) with a 150 µm polyacrylamide (PAA) gel (shown in teal) casting donut shaped cavity, is mounted on the glass in a 35mm glass bottom dish. pH-responsive gel (shown in yellow) is casted as a slab in the center of the donut shaped PAA gel. Cells of interest are cultured on top of the collagen-I coated PAA gel of defined stiffness. During GCM, the pH-responsive gel expands radially outwards and compresses the cells-PAA gel interface in turn, which is imaged in live (depicted by the imaging region of interest (ROI)). **G.** Representative image panels for 4 kPa PAA gel control, WT cells or M cells cultured on 4 kPa PAA gels for GCM. Pre-compression ROIs are shown to depict PAA gel alone or cells-PAA gel interface (marked by yellow dotted lines) as well as their respective post-compression ROIs representing the respective interfaces after 90 mins of continuous compression (marked by white dotted lines), Scale bar, 50 µm. **H.** Creep analysis where compressive strain (as fraction to the pre-compression perpendicular shift of the interface) incurred during GCM is plotted as a function of the corresponding time of compression in minutes, Data are mean±sem from three independent experiments. **I.** Analytical representation depicting the methodology and equations used to quantitate compressive strain and relative compressibility (relative to the respective PAA gel alone control). **J.** Relative compressibility measurement based on **I**, shows a distinct and lower value for WT population whereas M population shows no distinction compared to the PAA gel case. Data are mean±sem from three-four independent experiments. Statistical significance is calculated using Unpaired t-test with Welch’s correction. **K.** Representative image panels for 0.87 kPa PAA gel control, WT cells or M cells cultured on 0.87 kPa PAA gels for GCM. Pre-compression ROIs are shown to depict PAA gel alone or cells-PAA gel interface (marked by yellow dotted lines) as well as their respective post-compression ROIs representing the respective interfaces after 90 mins of continuous compression (marked by white dotted lines), Scale bar, 50 µm. **L.** Relative compressibility measurement based on **I**, shows a distinct and lower value for WT population as also seen in **J**. Importantly, M population shows a distinctly lower value compared to the PAA gel alone case but higher than the WT population. Data are mean±sem from three-four independent experiments. Statistical significance was calculated using Unpaired t-test with Welch’s correction.

where we have defined the ratio of cell area elasticities *k* = *K^M^_A_/K^WT^_A_*.

Based on our experimental findings that wildtype cells maintained their cell-cell junctions and established a durable, quiescent epithelial layer under ordinary circumstances (Figs. 3A,C,E,G,I and Supplementary Figs. 3A-B,E-G), we assigned a *P^WT^*_0_ value of 3.65 to these cells, which corresponded generically to a rigid jammed cells [33–35]. Simultaneously, the observation from experiments that HRas^V12^-expressing transformed cells exhibited less stable cell-cell junctions than wildtype cells (Figs. 3B,D,F,H,J and Supplementary Figs. 3C-G) implies that the transformed cells should have a larger value of 𝑃_0_, corresponding to softer unjammed cells. Furthermore, the larger homeostatic regions observed in transformed cells (Figs. 1E-F) implied a larger value of 𝐴_0_. Taken together, we propose that the competition of cellular junctional properties between M and WT cells can be encoded by a dimensionless cell shape index 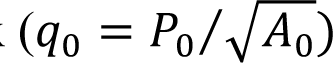. We first investigated the mechanical properties of the model cell layer by computing the hydrostatic pressure (𝛱) and interaction normal stress (𝜎_𝑛_) for the transformed and wildtype populations and the mechanical tension (𝑇_𝑖_) at cell-cell junctions[36] (Eq. SI-3-5 in SI Section). We observed that regardless of the size of the circular island, transformed cells exhibited lower junctional tension than wildtype cells when 𝑞^𝑀^_0_ > *q^WT^*_0_ (Fig. 4A and Supplementary Fig. 4A). Meanwhile, transformed cells experienced higher homeostatic pressure than wildtype cells when *A^M^*_0_ > *A*^*WT*^_0_ and *P^M^*_0_ > *P^WT^*_0_ (Fig. 4B). Further, we found that wildtype cells exerted compressive stress on transformed cells (𝜎_𝑀_ < 𝜎_𝑊_) (Supplementary Fig. 4B), with the stress gradient consistently oriented from high in wildtype cells to low in transformed cells. These phenotypic differences at the individual cell level provide the wildtype cells with a significant advantage in exerting compressive stress on the transformed population.

Consistent with the findings from our experiments (Figs. 1E-F), the model was able to accurately capture the density compaction observed in the transformed population within a competitive environment. Specifically, using the same compaction parameter 𝛥 (Eq. SI-6 in SI Section), which exhibited positive values when transformed cells were under compressive stress or elevated pressure (Fig. 4C), the simulations revealed an interesting finding that the increased compaction observed in transformed population is primarily influenced by the ratio of single cell area elasticities 𝑘 = *K^M^_A_/K^WT^_A_* (Fig. 4D). To quantify the effect, we calculated 𝐴 𝐴 the compressibility, C = 𝜕Δ⁄𝜕Π (carried out in the linear response regime, see SI Section for details). From the results shown in Fig 4D, we observed that when is 𝑘 < 1, the transformed cells exhibit greater compressibility compared to the wildtype cells, whereas for 𝑘 > 1, the compressibility of the wildtype cells exceeds that of the transformed cells. This provides compelling evidence for the crucial role of the elastic properties of tissue in determining its susceptibility to elimination by cell competition, which originates from mechanical imbalances at the microscopic cellular level.

### Gel compression microscopy (GCM) to measure collective compressibility

We then assessed whether the wildtype and transformed populations indeed differed in compressibility, as predicted by our theoretical modeling. To be able to reliably measure collective compressibility, we invented a novel biophysical technique termed as Gel Compression Microscopy (Figs. 4E-F and Supplementary Figs. 5A-B; for details see methods) or GCM.

Existing methods to study epithelial tissue mechanics rely either on atomic force microscopy-based indentation [37, 38], which only measures individual cell cortical stiffness but fails to sample bulk tissue mechanical properties, or on free-hanging monolayer compression, which requires cell-cell junction fidelity to sustain a suspended monolayer of cells [39, 40]. As we had observed previously, HRas^V12^-expressing cells have weak cell-cell junctions. Hence, we ruled out the free-hanging monolayer setup. We were, therefore, motivated to devise a unique technique where one can leverage live-imaging a compressing monolayer of cells in real-time (Figs. 4F-H) and quantitate its compressibility at the multicellular scale (Fig. 4I), while the given monolayer rests on a physiologically relevant substrate (Supplementary Figs. 5A-B).

The gel compression microscopy set-up consists of a cell growth-conducive polyacrylamide hydrogel of defined stiffness interfacing with an acrylic acid-based pH-responsive hydrogel (Supplementary Fig. 5A) [41]. The pH-responsive hydrogel expands at a buffer pH above 6 and contracts at a buffer pH below 4 [41]. For better control of the separation between the polyacrylamide gel and the pH-responsive gel and for reproducibility, we utilized soft lithography (Fig. 4E and Supplementary Fig. 5B) to micropattern molds of defined shape and size. Next, to carry out compressibility measurements, we cultured wildtype or HRas^V12^-expressing transformed cells to confluency on top of the polyacrylamide gel and subsequently immerse the interfacing pH-responsive gel with normal growth medium (pH ∼ 7.4) (Supplementary Fig. 5A).

Henceforth, the pH-responsive gel expands at a constant rate exerting a constant multicell-scale compressive stress on the wildtype or HRas^V12^-expressing monolayer grown over the polyacrylamide gel (Fig. 4H). Although, the empirical volume of substrate is much larger than the cells resting atop, we reasoned that any difference in cell behavior we might see between wildtype and transformed mutants will be comparable in the broad paradigm of the definitive parameters of our setup, Therefore, we computed relative compressibility of transformed monolayer and wildtype monolayer relative to that of polyacrylamide substrate alone (Fig. 4I). In our initial experiments, we used the polyacrylamide gel with Young’s modulus of 4 kPa, which is a physiologically relevant tissue stiffness [16]. On 4 kPa gel, the wildtype cell monolayer showed a significant and distinctly lower compressibility modulus relative to gel alone (Figs. 4G,J and Supplementary videos 2, 3), as indicated by our calculations of the relative compressibility modulus of 1.00±0.17 for gel alone compared to 0.49±0.07 for wildtype cell monolayer culture on the gel (Supplementary Fig. 5D). In contrast, the transformed cell monolayer with a relative compressibility modulus of 0.99±0.08 showed no difference in compressibility as compared to the gel-alone control (Fig. 4G,J, Supplementary Fig. 5D, and Supplementary video 4).

Based on these assessments, we wondered whether transformed cells possess a distinct but relatively smaller compressibility modulus than a 4 kPa gel. To this end, we performed the compressibility experiments with a softer polyacrylamide gel with Young’s modulus of 0.87 kPa (Fig. 4K and Supplementary videos 5, 6, 7). Here, we found that the transformed monolayer showed a lower relative compressibility modulus of 0.63±0.05 compared to the gel alone scenario of 1.00±0.13 (Fig. 4L, Supplementary Fig. 5D), pointing towards a small but distinct mechanical elasticity contributed by the transformed population. Interestingly, whilst transformed monolayer is less compressible than the soft gel but, at least, it still is relatively two-fold more compressible than the wildtype monolayer, which amounts to a relative compressibility modulus of 0.31±0.03 (Fig. 4L, Supplementary Fig. 5D).

Taken together, as had been predicted by our simulations (Fig. 4D), the compressibility measurements with gel compression microscopy suggested that there exists a distinctively higher collective compressibility of the transformed population as compared to their wildtype counterpart and that in line with our predictions, forms the basis of competition driven compressive stress during epithelial defense against cancer.

### Different regimes of mechanical phenotypes in cell competition

Our experiments on active compression revealed that the transformed mutant (M) cells exhibit higher collective tissue compressibility than their wildtype (WT) counterparts (Figs. 4J,L), consistent with our previous linear-response measurements (Fig. 4D). This difference in collective tissue elasticity between the competing populations represents a key mechanical cue driving non-proliferative cell competition. To further understand the underlying factors that contribute to this competition, we incorporated active compression into the SPV. This was achieved by allowing the wildtype cells to exert an active self-propelled force on the transformed island (Eq. SI-7,8, see SI Section for a detailed discussion), thereby generating collective elastic response. When an active force was applied, the transformed cells experienced compression. By quantifying the relative change in the total area occupied by the transformed cells, we derived the compressive areal strain using Eq. SI-9. This enabled us to calculate the bulk modulus B (Eq. SI-10), which represents the material’s response to isotropic compressions (also done in the linear response regime, see SI Section for a detailed discussion). In essence, the bulk modulus of a material is inversely related to its compressibility. We measured B for a small magnitude of the active force (see SI Section) and a wide range of values of 𝑘, *A^M^*_0_ and 𝛿𝑞_0_ = 𝑞^𝑀^ − *q^WT^*_0_ with *q^WT^*_0_ = 3.65 as reported in the SPV modeling section of this work. Fig. 5A illustrates that the differential bulk modulus (𝛥 = 𝛥_𝑀_ − 𝛥_𝑊_) between wildtype and transformed cells are controlled by both 𝑘 and 𝛿𝑞_0_. Notably, 𝛥 increases linearly with 𝑘 *A^M^*_0_, where the slope is independent of 𝛿𝑞_0_ (Fig. 5B). After subtracting by an offset that depends on 𝛿𝑞_0_, the data collapsed (Fig. 5B inset, see SI Section for details) to a simpler functional relationship between 𝛥, 𝑘 *A^M^*_0_ and 𝛿𝑞_0_, given by:

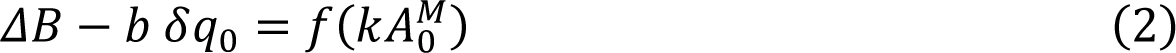

**Figure 5.**
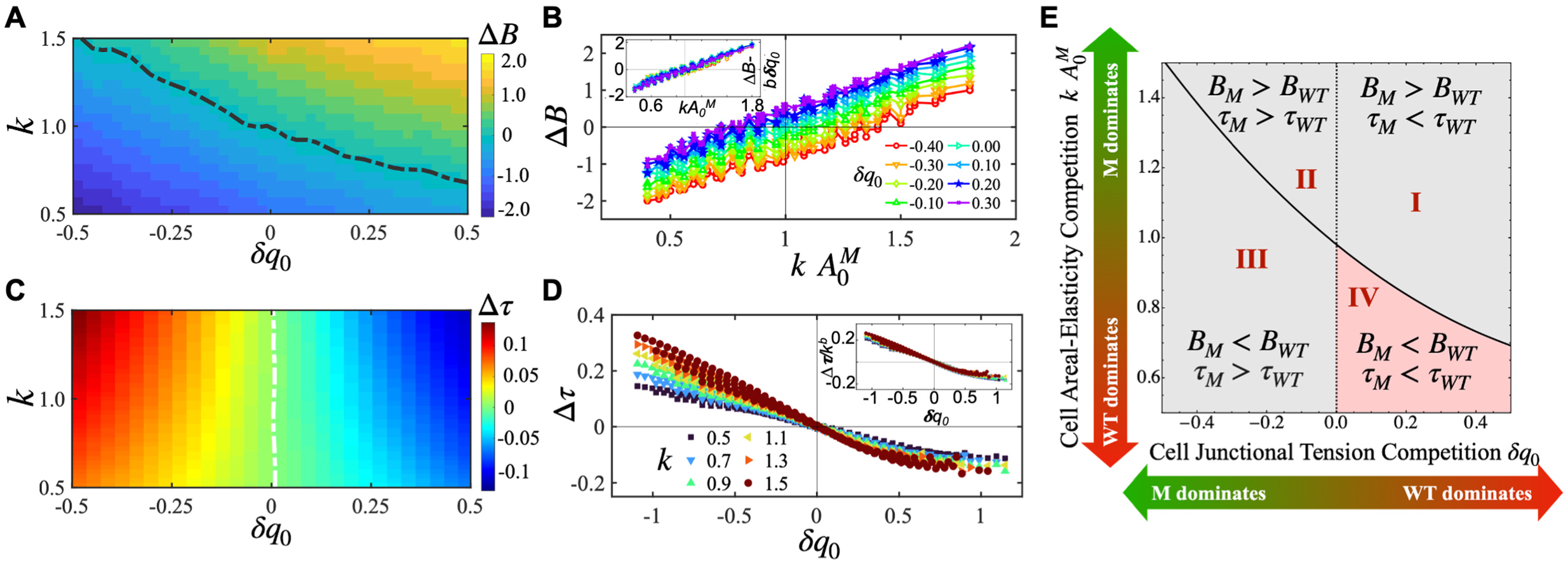
Theoretical modeling reveals different regimes of mechanical phenotypes in cell competition. In the computational simulations, we assessed both the bulk and shear moduli of the cell colonies. The bulk modulus provided insights into a tissue’s mechanical response to isotropic compression, while the shear modulus was essential for evaluating the tissue’s reaction to shear forces. These mechanical properties were analyzed for a wide range of parameters 𝑘, *A^M^*_0_ and 𝛿𝑞_0_. Here, 𝑘 encoded competition in cell area elasticities between the cell types. When 𝑘 < 1, the transformed mutant (M) colony was more compressible than the wildtype (WT) cells surrounding them. On the other hand, 𝛿𝑞_0_encoded the competition in cellular junctional properties; when 𝛿𝑞_0_ > 0, the transformed cells had weakened cellular junctions compared to the WT cells. **A.** Behavior of the differential bulk modulus 𝛥 between WT and M cells in the 𝑘 − 𝛿𝑞_0_ plane for *A^M^*_0_ = *A*^*WT*^_0_ = 1. The black dotted line corresponds to 𝛥 = 0. **B.** 𝛥 0 0 plotted as a function of 𝑘 *A^M^*_0_ for a range of 𝛿𝑞_0_ values. The inset shows 𝛥 − 𝑏 𝛿𝑞_0_ as a function of 𝑘 *A^M^*_0_ for the same data. Here we used 𝑏 = 1.7 ± 0.2. **C.** The behavior of the differential colony tension 𝛥𝜏 in the 𝑘 − 𝛿𝑞_0_ plane for *A^M^*_0_= 1. The white dotted line corresponds to 𝛥𝜏 = 0. **D.** 𝛥𝜏 plotted as a function of 𝛿𝑞_0_ for a range of 𝑘 values (and *A^M^*_0_ = 1). The inset shows 𝛥𝜏/𝑘^𝑑𝑑^ as a function of 𝛿𝑞_0_ exhibiting a scaling collapse. Here we find 𝑑 = 0.53 ± 0.12. Both the scaling parameters in **B** and **D** are obtained by minimizing error functions which we defined to measure the closeness of the respective data sets. **E.** based on the analysis of Fig. 5A**-D**, we predict a phase diagram which qualitatively distinguishes between mechanical phenotypes that are favored in non-proliferative mechanical cell competition. Specifically, a colony of cells with lower 𝛿𝑞_0_and higher 𝑘 *A^M^*_0_ will have a competitive advantage, as they exhibit lower compressibility and higher rigidity. The predicted phase diagram serves as a useful tool for identifying such mechanical phenotypes.

These results indicate that competition in *cell area elasticity* plays a central role in driving the differences in bulk modulus between M and WT cells. Conversely, the parameter 𝛿𝑞_0_, which represents competition in cellular junctional properties, exerts a more subordinate influence.

While the bulk modulus provides insights into a tissue’s mechanical response to isotropic compression, the shear modulus is essential for evaluating the tissue’s reaction to shear forces. Notably, both solid and fluid materials possess a bulk modulus, whereas the shear modulus is a characteristic exclusive to solids. As a material transitions from a solid state to a flow state, its shear modulus diminishes to zero. Here instead of calculating the shear modulus explicitly for M and WT colonies, we used the mean edge tension (Eq. SI-11) as an approximation for the shear modulus (see SI Section for details). Previous research has demonstrated a linear relation between these two quantities [42].

In Fig. 5C-D, we observe that the differential tension (Δ𝜏 = 𝜏_𝑀_ − 𝜏_𝑊_) decreases as 𝛿𝑞_0_ increases, this occurs for all values of 𝑘, which only sets the rate of decrease. Fig. 5D inset demonstrates that the functional relationship between Δ𝜏, 𝛿𝑞_0_, and 𝑘 for *A^M^*_0_ = 1 exhibits a universal scaling collapse (see SI Section for details) described by:

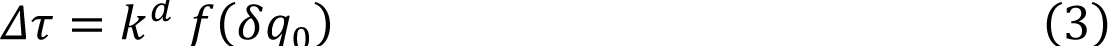

These results indicate that the competition in cellular junctional properties (𝛿𝑞_0_) plays a central role in driving the differences in shear modulus between M and WT cells. Conversely, competition in cell area elasticity (𝑘) exerts a more secondary influence.

Our simulations show that, in addition to differences in homeostatic cell area (𝐴_0_), cells can compete via differences in their area elasticity (𝑘) and junctional properties (𝛿𝑞_0_). Taken together, in Fig. 5E we propose a phase diagram that identifies four possible regimes of mechanical competition (see SI Section for details). Given that transformed cells exhibit weaker cellular junctions than their wild-type counterparts, our theoretical model predicts a reduced shear modulus for these cells. Simultaneously, their increased compressibility results in lower cell area elasticity, leading to a diminished bulk modulus compared to WT cells. These findings lead to the conjecture that successful elimination of transformed cells by cell competition requires the competing populations to emerge as winners in Region II and losers in Region IV (shaded red in Fig. 5E) of this mechano-phenotypic phase diagram.

### Role of destabilized cell-cell junctions in transformed cells

Finally, we attempted to understand the mechanistic basis for this difference in collective compressibility between the transformed and normal cell populations. At the single-cell level, atomic force microscopy experiments have delineated that HRas^V12^-expressing transformed cells are stiffer than wildtype cells when in an adherent state, whilst they are softer when in loosely attached or suspended form [37, 38, 43]. In contrast, we found higher collective compressibility in transformed cells. One possible explanation for these two apparently discrepant observations is the weaker cell-cell adhesions in the transformed cells as compared to the wildtype cells. Indeed, junctional E-cadherin mislocalization leading up to weaker cell-cell adhesions, upon Ras transformation, has been shown before [44]. Therefore, it is possible that weak cell-cell adhesions make a collective of transformed cells more compressible than a collective of wildtype cells. Relevantly, it is known that a matrix metalloproteinase-dependent cleavage of the extracellular domain of the cell-cell adhesion protein E-cadherin drives epithelial extrusion [45].

We, therefore, hypothesized that cell-cell adhesions in the HRas^V12^-expressing mutants could be compromised due to the cleavage of the extracellular domain of E-cadherin. To test this hypothesis, we used DECMA-1 antibody which binds to the extracellular domain of E-cadherin. Upon mild detergent pretreatment, which permeabilizes cell membrane as well as partially disrupts the interaction between two E-cadherin molecules from the adjacent cells, DECMA-1 detects both cytoplasmic and membrane-bound E-cadherin, provided their extracellular domains are intact. Without the detergent, DECMA-1 detects only those E-cadherin molecules that are not bound to other E-cadherin molecules and have their extracellular domains intact. We stained both wildtype and transformed cells with this antibody with or without the detergent pretreatment. With detergent pretreatment, we found that E-cadherin remains intact at the junctions of the transformed cells and wildtype cells (Fig. 6A). However, without membrane permeabilization, DECMA-1 showed staining only at the junctions of the transformed cells (Fig. 6B). These results revealed that E-cadherin molecules in transformed mutants remains intact but not bound to E-cadherin molecules of the adjacent cells.

**Figure 6.**
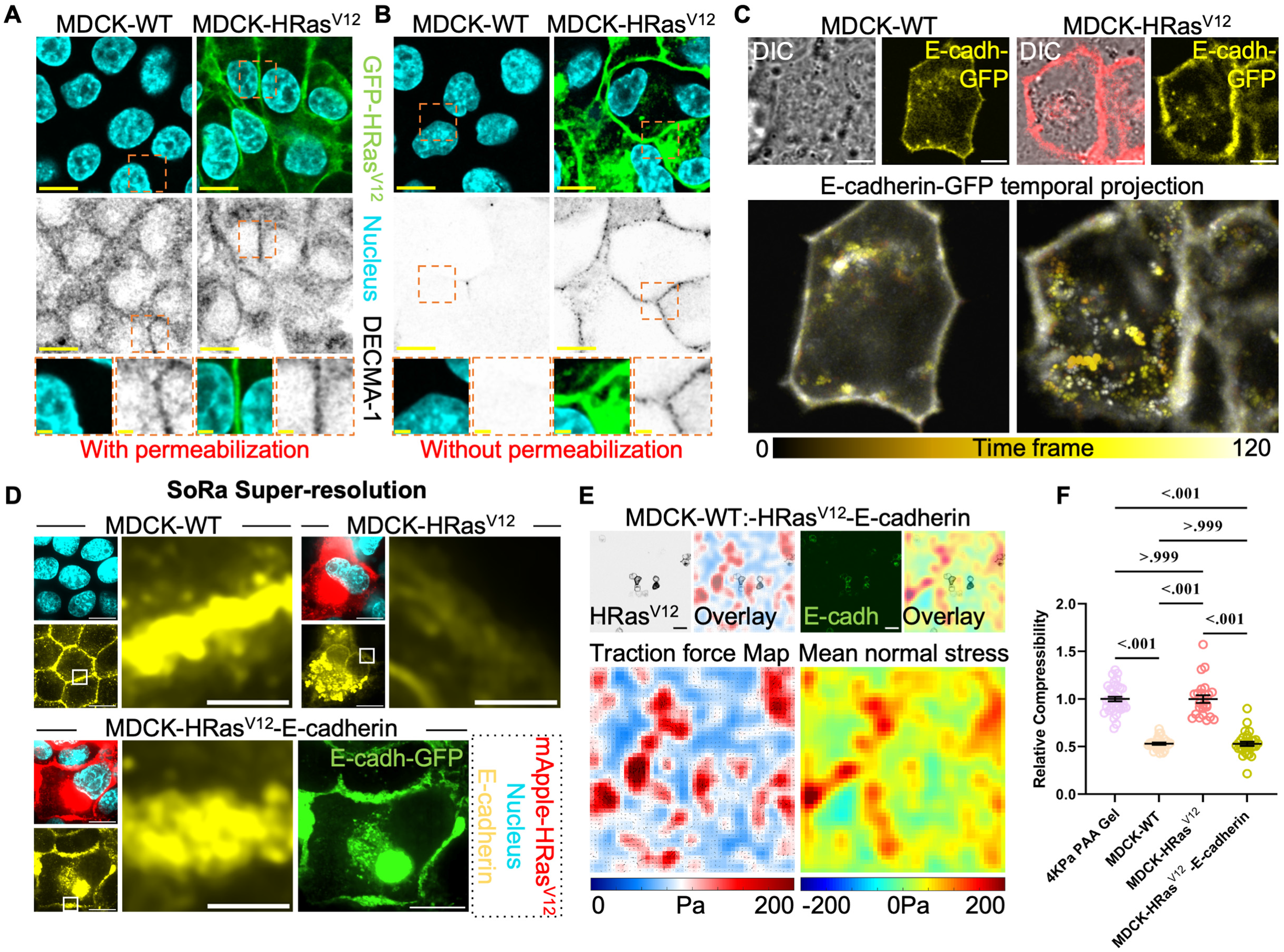
Compromised E-cadherin based cell-cell junctions in the HRas^V12^-transformed mutants underlie competition induced compressive stress as well as rescued collective compressibility. **A,B.** Representative image panels to show immunostained MDCK-wildtype (-WT) or MDCK-HRas^V12^-transformed mutant cells for E-cadherin extracellular domain marked by DECMA-1. The immunostaining is performed A, WITH detergent (TritonX-100) or **B**, WITHOUT detergent to demonstrate the inherent instability in MT-MT junctions but not WT-WT junctions whereas the extracellular domain being intact in either case. Scale bar, 10 µm. **C.** E-cadherin live dynamics are studied in WT or MT cells mildly overexpressing E-cadherin-GFP. Representative differential interference contrast (DIC) and E-cadherin-GFP image panels are shown. Temporal dynamics for each cell-type is shown as a color projection map where each time stamp is differentially colored in the ‘yellow hot’ color panel and superimposed. Each time frame refers to 5 second imaging window. Scale bar, 5 µm. **D.** Super-resolution spinning-disk confocal microscopy using optical photon reassignment (SoRa) image panels to show endogenous E-cadherin in WT, MT or MT overexpressing E-cadherin-GFP. Junctional E-cadherin is shown in insets from the three scenarios to depict E-cadherin loss in transformed cells and rescue in transformed cells overexpressing exogenous E-cadherin-GFP. Scale bar, 10 µm for all image panels and 2 µm for all insets. **E.** Traction force microscopy and stress microscopy panels for cell competition between WT cells and MT cells overexpressing E-cadherin-GFP. Competition induced compressive stress as seen in **Fig. 2C**, is evidently rescued when E-cadherin is supplied exogenously in the transformed cells. Scale bar, 10 µm. **F.** Relative compressibility measurement based on **Fig. 4I**, shows a distinct and lower value for WT population whereas MT population shows no distinction compared to the polyacrylamide gel control, as also seen in **Fig. 4J**. Importantly, MT cells overexpressing E-cadherin-GFP show a distinctly similar value to the WT case, showcasing genetic rescue to the mechanical phenotype of compressibility differential. Data are mean±sem from three-four independent experiments. Statistical significance was calculated using Unpaired t-test with Welch’s correction.

To understand the reason behind improper E-cadherin coupling, we then studied the dynamics of E-cadherin in live cells expressing a E-cadherin-GFP construct. We observed an enhanced transport of E-cadherin in transformed cells but not in wildtype cells (Fig. 6C and Supplementary video 8), where E-cadherin molecules showed distinct vesicular fingerprints. Interestingly, this enhanced transport was not due to the endocytic awakening [46] since none of the endocytic compartments, including Rab5 for early endosomes, Rab11 for recycling endosomes, Rab7 for late endosomes and LAMP1 for lysosomes, showed any preferential co-localization with E-cadherin in transformed cells (Supplementary Figs. 6A-E).

In addition, using SoRa super-resolution imaging, we had found that junctional E-cadherin significantly deteriorated in the junctions of transformed cells (Figs. 3B,D,F,H,J). To recover the junctional E-cadherin pool in the transformed cells, we overexpressed E-cadherin-GFP in transformed cells. We assumed that E-cadherin overexpression would lead to the recovery of junctional E-cadherin concentration, and because of increased concentration, it would also lead to enhanced coupling of E-cadherin molecules, effectively “gluing” the transformed cells. Super-resolution imaging with SoRa revealed that junctional E-cadherin pool was indeed recovered within wildtype range upon E-cadherin overexpression (Fig. 6D). Further, when we performed traction force microscopy coupled with stress microscopy in the competition setup with wildtype cells and these E-cadherin-GFP overexpressing cells, there appeared to be no compressive stress at the transformed loci (Fig. 6E) unlike what we had observed before with the transformed cells alone (Fig. 2C). In fact, just by enriching junctional E-cadherin pool, we could rescue the competition dependent compressive stress which the transformed mutants would experience otherwise, when surrounded by wildtype cells (Fig. 6E). Finally, we performed gel compression microscopy to evaluate the collective compressibility of the E-cadherin-GFP overexpressing HRas^V12^-transformed cells (Supplementary Fig. 7A and Supplementary video 9). Indeed, collective compressibility of these transformed mutants was now rescued to the range of wildtype population (Fig. 6F) and thus explains the rescue in compressive stress when these transformed cells encounter wildtype cells in their vicinity.

## Discussion

In this work, we combined experimental investigation at the wildtype-vs-transformed cell competition regime in the small intestinal epithelia of mice as well as in cultured MDCK epithelia, along with robust *in silico* modeling, to provide direct mechanobiological evidence towards epithelial homeostasis. We, therefore, attempted to define the mechanistic and mechanical basis of epithelial homeostasis mediated by cell competition. We present *in vivo* evidence to define transformation-induced mechano-phenotypes pertaining to distinct biophysical aspects of cells once they incur HRas^V12^ mutation load. These phenotypes include higher preferred homoeostatic area, and compromised adherens-based cell-cell junctions, which are characterized also by transformed MDCK cells in culture. Further, we discovered pre-elimination compressive stress, which is unique to cell competition and henceforth, may be called <u>c</u>ompetition-induced <u>c</u>ompressive <u>s</u>tress (CICS).

Our work interestingly shows that a growth rate difference between the competing populations, which is classically considered to be the driver of cell competition and oncogenesis, does not seem to be a critical factor for cell competition against HRas^V12^-expressing transformed cells. Instead, we hypothesize that, the general mechanobiological aspects pertaining to epithelial cells which we show to be perturbed upon transformation, might be the drivers of competition. To reconcile these observations in a single framework, we utilized a computational Self-Propelled Voronoi model and show that a collective compressibility differential might explain the emergence of compressive stress during cell competition. While the existing tools to quantitate collective compressibility are either too complex, too customized to be implemented, or rely entirely on the impending cellular strength to suspend freely, we developed an inexpensive, physiologically compatible technique called gel compression microscopy (GCM). GCM includes a compliant substrate for monolayer adhesion, whose stiffness can be modulated, and enables real-time measurements of collective compressibility. Using GCM, we showed that the transformed cell population is indeed more compliant to in-plane compression. GCM measurements thus revealed an oncogenic transformation in the epithelial cells leading to a multicellular length-scale change in its material property. Taken together, we present a mechanical picture where the oncogenic transformation of a group of cells in an otherwise normal epithelium modulates their cellular biophysical properties making them more compressible as a collective. Subsequently, during cell competition, a local compressive stress emerges at the transformed loci, which leads to a successful epithelial homeostasis and possibly, a fundamental defense against cancer.

In this regard, a recent study parameterized differences in cellular-scale measurements during cell competition between wildtype and transformed cells lacking a tumor suppressor protein, Scribble [47]. They used an agent-based model to show that the homeostatic density of the competing cell types primarily governs mechanical cell competition, while the local tissue organization influences the outcome and kinetics of biochemical competition [47]. However, in the absence of direct measurement of forces, a comprehensive understanding of the causal mechanism driving the mechanical cell competition remained missing. In contrast, we first used stress microscopy techniques to discover CICS. Subsequently, our efforts to understand the mechanistic origin of CICS led to the discovery of differential compressibility via computational modeling and the invention of GCM. Hence, our study went much beyond the previous geometric representation of cell competition and elucidated how individual cell mechanics integrate at the multi-cell scale to bring about oncogenic cell elimination through competition. Together, it is tempting to speculate that the transformed cells struggle to achieve their desired, higher cellular area while being resisted by the surrounding normal cells, giving rise to CICS.

In the context of stable epithelial monolayers both *in vitro* and *in vivo*, the factors that trigger cell competition, including the Ras family of proteins, Myc, Toll, and apicobasal polarity proteins, are also involved in modulating junctional tension [14, 48–50]. Consistent with these observations, we demonstrate that junctional fidelity in HRas^V12^-expressing transformed cells is indeed aberrant in culture and physiology. The most important epithelial cell-cell adhesion protein, E-cadherin, loses its junctional localization upon oncogenic transformation (Fig. 3). The membrane-anchored E-cadherin molecules get actively displaced from the junctions of the transformed cells, trafficked through the cytoplasm and are not turned over back to the junctions in time, compromising junctional stability (Fig. 6C, Supplementary Fig. 6 and Supplementary video 8). This instability is corroborated by the diffused cytoplasmic occurrence of another protein associated with cell-cell junctional maturation, alpha-catenin (Supplementary Fig. 3). Remarkably, just by enriching the overall pool of available E-cadherin in the mutants, we observed a significant recovery in their collective compressibility, and the CICS disappeared (Figs. 6E-F).

In the paradigm of cell competition, what still remains to be explored is the cellular phenotypes that fundamentally perturb the bulk mechanical elasticity of epithelial cells as they undergo transformation. How cell-autonomous and non-cell-autonomous features arise in normal as well as transformed cells to guide mechanobiological patterns of cell competition, remains poorly understood. Finally, it will be interesting to investigate whether compressive stress also appears during competition against mutations in other Ras family proteins and whether the underlying mechanism can be generalized beyond HRas^V12^. Nevertheless, using physiologically relevant model systems, our comprehensive study has unveiled the previously enigmatic physical principle and molecular mechanism that underlie cell competition in epithelial homeostasis. This breakthrough not only provides crucial insights into the fundamental dynamics of this process but also opens exciting avenues for the emerging field of mechano-medicine. By understanding how mechanical forces and tissue compressibility play a pivotal role in cell competition, we have set the stage for innovative therapeutic strategies that leverage these principles to target and potentially mitigate the spread of cancerous cells within tissues.

## Materials and Methods

### Mice work and tissue sectioning

All animal experiments were conducted under the guidelines set by the Animal Care Committee of Kyoto University. C57BL/6 mice of either sex were used in this study. villin-CreER^T2^ mice were crossed with DNMT1-CAG-loxP-STOP-loxP-HRas^V12^-IRES-eGFP mice to generate villin-HRas^V12^-GFP [12]. PCR genotyping of mice was performed using primers listed in Supplementary Table 5. Mice heterozygous for each transgene were used for experiments. 6-12-week-old villin-HRas^V12^-GFP mice were given a single intraperitoneal injection of 2.0 mg of tamoxifen (Sigma-Aldrich) in corn oil (Sigma-Aldrich). These mice were sacrificed 2 days after tamoxifen injection and perfused with 1% paraformaldehyde (PFA) (Sigma-Aldrich) in phosphate-buffered saline (PBS). The isolated tissues were fixed with 1% PFA in PBS for 24 hours and incubated in 10% sucrose in PBS for 6 hours, followed by a 1-day incubation in 20% sucrose in PBS at 4°C. The tissues were embedded in Tissue-Tek optimal cutting temperature compound (SAKURA) and 10-μm-thick frozen sections were cut using a cryostat (Leica) and mounted on glass slides (Matsunami). The sections were incubated with 1× Block-Ace (DS Pharma Biomedical) and 0.1% Triton X-100, 1% Normal Donkey Serum (NDS) and 1% Normal Goat Serum (NGS) in PBS for 1 h, followed by incubation with primary or secondary antibody diluted in PBS containing 0.1× Block-Ace, 0.1% NDS, 0.1% NGS and 0.1% Triton X-100 for 2 h or 1 h, respectively, at room temperature. The stained samples were covered with mounting media (VectaMount, Vector labs), mounted gently with glass coverslips (Matsunami) and dried overnight before proceeding to microscopy.

### Cell culture

Madin–Darby canine kidney (MDCK) epithelial cells were used in this study. Tetracycline-resistant wild-type MDCK (MDCK-WT) line and tetracycline-inducible HRas^V12^-expressing MDCK (MDCK-GFP-HRas^V12^). MDCK cells were cultured in Dulbecco’s modified Eagle’s medium (DMEM, Gibco) supplemented with GlutaMax (Gibco) and 5% fetal bovine serum (tetracycline-free FBS, Takara Bio) along with 10 U ml^−1^ penicillin and 10 μg ml^−1^ streptomycin (Pen-Strep, Invitrogen). Cells were maintained at 37 °C and 5% CO_2_ unless mentioned otherwise. To set up cell competition, a mosaic monolayer constituting normal MDCK cells (MDCK-WT) and transformed MDCK cells (MDCK-GFP-HRas^V12^) were cultured overnight in a ratio (40:1, respectively) on collagen-I (Invitrogen, A1048301) coated polyacrylamide (PAA) hydrogels of varying stiffness in the absence of tetracycline (Supplementary Fig. 2A). Cell competition was induced only after a confluent monolayer of MDCK cells was formed. Thereafter, GFP-HRas^V12^ expression was induced by supplementing 10 μg ml^−1^ a tetracycline derivative, doxycycline (Sigma-Aldrich) to the culture medium.

MDCK-WT cells expressing tetracycline-inducible mApple-HRas^V12^ (MDCK-mApple-HRas^V12^) were created in this work. To establish stable cell lines, MDCK cells were transfected with respective plasmid DNA using Lipofectamine 2000 (Invitrogen, 11668019). Appropriate selection for positives was done by supplementing culture medium with 800 μg ml^−1^ Geneticin (Invitrogen, 10131035) or 500 μg ml^−1^ Zeocin (Invitrogen, R25001). Transient transfection with plasmids was done using Lipofectamine 2000 following manufacturer’s protocol. Post 8–12 h of transfection, cells were trypsinized and seeded onto hydrogel substrates and cultured overnight. Once confluent, cells were either fixed and immuno-stained or imaged directly.

### Polyacrylamide (PAA) Hydrogel preparation for compliant ECM

Cells were cultured on compliant hydrogel substrates for the presented studies, where polyacrylamide hydrogels functionalized with collagen-I were generated as described previously [51, 52]. 4% (3-Aminopropyl) triethoxysilane (APTES) (Sigma-Aldrich)-treated and 2% glutaraldehyde (Sigma-Aldrich)-activated 35mm glass-bottom dishes (Ibidi) were used to cast thin (120 μm – 150 μm) polyacrylamide (PAA) hydrogel substrates. Hydrogel substrate of 4kPa elastic modulus were used for all the experiments unless stated otherwise. For Gel Compression Microscopy experiments, hydrogels of elastic modulus 4kPa and 0.87kPa were used. The hydrogel substrates of varying elastic modulus were prepared by mixing the desired volume of 40% acrylamide (Sigma-Aldrich) and 2% bisacrylamide (Sigma-Aldrich) as given in Supplementary Table 1. Gel surfaces were functionalized with sulfosuccinimidyl-6-(4′-azido-2′-nitrophenylamino) hexanoate (Sulfo-SANPAH, ThermoFisher 22589) and coated overnight with collagen-I at 4 °C to ensure cell attachment. Cells were seeded onto the gel area and grown until confluency.

### Immunofluorescence

Cells cultured on polyacrylamide hydrogels were fixed with 4% formaldehyde (ThermoFisher) diluted in 1X Phosphate buffered saline (PBS, Sigma-Aldrich) for 15 minutes at room temperature. After washing away the fixative with 1X PBS, cells were treated with blocking/staining solution (0.3% TritonX-100 (Sigma-Aldrich) in 1X PBS supplemented with 2% Bovine Serum Albumin (BSA, Himedia)) for 1 hour at room temperature. Further, cells were incubated with primary antibody prepared in blocking/staining solution overnight at 4°C. Post-primary antibodies incubation, cells were washed thrice with 1X PBS and incubated with secondary antibodies conjugated to Alexa fluor 568 or 647 and 4′,6-diamidino-2-phenylindole (DAPI) to mark the cell nucleus, prepared in blocking/staining solution, for 1 hour at room temperature. Finally, the samples were washed three times with 1X PBS before proceeding to microscopy.

### Antibodies and Plasmids

Source and dilution information for all primary and secondary antibodies used in immunofluorescence staining are given in Supplementary Table 2. Details of plasmids used in this study are listed in Supplementary Table 3.

### EdU labelling

EdU incorporates into dividing cells and therefore quantification of EdU^+^ cells can be used as a direct measure of cell proliferation [53]. For EdU labelling, cells were pulsed with 10 μM EdU for 1h and fixed with 4% paraformaldehyde for 10-15 min at room temperature. the incorporated EdU detected using a Click-iT EdU imaging kit (ThermoFisher Scientific) according to the manufacturer’s instructions.

### Traction force and monolayer stress microscopy

Traction force microscopy was performed as described previously [54]. Briefly, glutaraldehyde-activated glass bottom dishes (Ibidi) were used to cast thin polyacrylamide (PAA) gel substrates (Young’s modulus of 4 or 11 kPa) containing 0.5 μm fluorescent carboxylated-polystyrene beads (Sigma-Aldrich). These gel surfaces were then functionalized with Sulfo-SANPAH and coated with 300 µg ml^−1^ collagen-I to ensure cell attachment. Cells were seeded in the confined areas and grown until a confluent monolayer was obtained. Subsequently, confinement was released by lifting off the PDMS block, and images for cells and beads were acquired as the cells migrated. After the experiment, cells were trypsinized, and resulting bead positions in a relaxed state were obtained (i.e., reference images). The displaced images were aligned to correct for drift and compared to the reference image using particle image velocimetry to create a regular field of displacement vectors with a grid spacing of 5.44 μm. Displacement vectors were interpolated using cubic splines. From these vectors, traction stresses were reconstructed using regularized Fourier Transform Traction Cytometry [55] with a regularization parameter chosen by Generalized cross-validation [56]. Stresses within the monolayer were then calculated by Bayesian inversion stress microscopy (BISM) from the cell-substrate tractions using a force balance algorithm written in MATLAB (MathWorks) as described previously [57].

### Gel Compression Microscopy

Gel compression microscopy (GCM) is an inexpensive and novel biophysical technique described in this work, which utilizes a monotonically expanding pH-responsive gel to compress a polyacrylamide (PAA) gel at its interface. In doing so, GCM offers to study and quantify the compressibility and related mechanical properties of the PAA gel (and the cell monolayer cultured on top). Furthermore, since the technique involves live cell collectives, it allows for capturing the evolution of compressive stress in real time. This section elaborates on the conceptual as well as technical aspects of GCM, which are illustrated in Fig. 4 and Supplementary Fig. 5.

#### pH-responsive gel preparation

pH-responsive hydrogel was prepared as described before [41]. Briefly, the pH-responsive gel mix was prepared as weight ratios of the chemical species, all of which are mentioned in Supplementary Table 4. The mix was sonicated briefly to homogenize and vortexed to completely dissolve all the components together. Further, the prepolymer mix was stored under refrigeration and protected from light. To create pH-responsive gel slabs for GCM, prepolymer mix was poured into 6mm-diameter pillar holes in circular Polydimethylsiloxane (PDMS) stamps. The mix was then cured under the highest power setting of the Analytik Jena UVP crosslinker CL-1000 for 90 minutes. The polymerized pH-responsive hydrogel slabs were washed once with milli-Q water to remove uncured polymer and further stored in 1xPBS (pH-2.5) until required.

#### PAA gel preparation

To cast PAA gels of defined stiffness for GCM experiments, first, the PDMS stamps were taken out of isopropanol onto a dry kimwipe, pat-dried and fully dried in a hot air oven set at 72°C for 1 hour. The stamps were then mounted ‘pillar side down’ on the glass coverslip of a 35mm glass bottom dish (Ibidi) in order to leave a donut-shaped gap of defined height (∼150 um) between PDMS and glass. This setup was then moved into a degassing chamber (Tarsons) which was connected to a Vacuum-Argon dual line. Three sequential rounds of vacuuming (15 mins at high pressure until the chamber pressure reaches 0.2-0.5 torr) and gassing with argon (until the chamber pressure is relieved). Finally, under constant argon flow, a prepared PAA gel mix (referred to in Supplementary table 1) was injected into the gap between PDMS and glass. The setup is left in the closed gas chamber under constant argon flow for 1 hour to allow ample time for PAA gel polymerization. After the polymerization step, PDMS stamps are gently peeled off from the glass. The polymerized donut-shaped gels (shown in Supplementary Fig. 5B) are hydrated with milli-Q water before proceeding for activation to make them compliant for cell seeding as mentioned in a previous section.

#### GCM setup

As a first step, cells of interest are cultured on collagen-I coated donut-shaped PAA gels until confluency. Cells are then washed once with 1xPBS and covered with excess of culture media. A prepared cylindrical pH-responsive of ∼2 mm height is placed in the center empty region of the donut-shaped PAA gel. The setup is left unperturbed on the bench for 5 minutes before placing a glass coverslip on top of the pH-responsive gel. A glass cylinder is further placed on top of the coverslip and holds the dish cap on top, to keep the pH-responsive gel from floating away. The entire setup is illustrated in Supplementary Figs. 5A-B. Once set up, the system is moved to a microscope and regions of interest to be live-imaged are marked (shown in Figs. 4G,K).

#### Compressibility analysis with GCM

The data acquired with GCM shows progressive compression of the cell monolayer/PAA gel in response to the radially outward expansion of the pH-responsive gel. An arbitrary line perpendicular to the cell monolayer/PAA gel drawn from the interface with pH-responsive gel extending 1000 μm deep into the monolayer is taken as the “initial length (L_x_)”. A progressive reduction in the length of this line is taken as “compressed length (L_x_’)” for that particular point in compression. The progression of ′ Compressive strain, 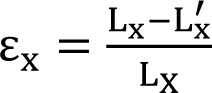, is plotted as a function of time to map the ‘creep’ the incurs x _LX_ during GCM. Furthermore, mapping the compressive strain over ‘pre-compression’ at the start of compression and ‘post-compression’ at the end of a given point in time and averaging over many such events gives a compressibility modulus which is compared between cell-types to judge their compressibility differences.

### Live cell imaging

For the live membrane dynamics imaging, live cell membranes were labelled with the CellMask (Invitrogen, C10046) probe coupled to AlexaFluor 647 based on manufacturer’s guidelines. For live E-cadherin dynamics, MDCK -wildtype or -transformed cells were transfected with E-cadherin-GFP plasmid (Addgene) using Lipofectamine 2000 transfection reagent (Invitrogen, 11668019) following manufacturer’s protocol. Moderately overexpressing GFP^+^ cells were chosen for these experiments. Time-lapse images of live samples were acquired in the Olympus FV3000 laser scanning setup using the supplied live-cell chamber maintained at 37°C. HEPES (Gibco) buffer at a final concentration of 50 μM was added to the culture medium. Live imaging for gel compression microscopy was done using ×20 objective (0.40 HC PL FLUOTAR L) on Leica DMI8 inverted microscope using Leica Application Suite (LasX v3.7.0.20979). Time-lapse images of live samples were acquired in the Leica DMI8 wide-field setup using a Pecon live-cell chamber maintained at 37°C and supplied with humidified 5% CO_2_.

### Microscopy

Fluorescence images were acquired using 60X water immersion objective (UPLSAPO W, N.A. = 1.20, Olympus) mounted on an Olympus IX83 inverted microscope equipped with a scanning laser confocal head (Olympus FV3000), Olympus FV31-SW (v2.3.1.198). For Super-Resolution via Optical Reassignment [29] or SoRa imaging, images were acquired with a x60 water immersion objective (Plan Apo VC 60XA, N.A =1.20, Nikon) mounted on a Nikon inverted microscope supplied with Yokogawa CSU-W1 (SoRa Disk) scanner. Images were further processed via the ‘clarify’ module in-built to the NIS elements imaging software (v5.42.01) provided with the system. Wide-field images to measure proliferation rates were acquired using ×20 objective (0.40 HC PL FLUOTAR L) on a Leica DMI8 inverted microscope using Leica Application Suite X (LAX, v3.7.0.20979).

### Soft photolithography-based micropatterning

Polydimethylsiloxane (PDMS) stencils with pillars of defined shapes were created using soft photolithography [58]. Briefly, 150 um thick SU-8 2075 (Kayaku advanced materials, Y111074) was coated on a silicon wafer followed by soft baking at 65° C for 5 minutes and 95° C for 20 minutes. SU-8 was exposed in the desired pattern using the MicroWriter ML 3 Pro (Durham Optics Magneto Ltd) followed by post-exposure baking at 65° C for 7 minutes and 95° C for 15 minutes. The unexposed photoresist was removed by immersing the wafers in the SU-8 developer (Kayaku advanced materials, Y020100) for 8-10 minutes. Patterned SU-8 wafers were then hard-baked at 170° C for 45 minutes. PDMS (Sylgard 184, Dow Corning) was mixed in a ratio of 1:10, degassed, and poured over the patterned wafers. After curing at 65° C for 4 h, stencils were cut out and were peeled off. Since polyacrylamide (PAA) preparations do not polymerize in a PDMS cast and there is no protocol available in the literature, we standardized a way of preparing PAA gels in a PDMS environment. First, the prepared PDMS stencils were dipped in excess absolute isopropanol and sonicated at 37 kHz for 1 hour in a sonicator bath (Fisherbrand, FB11203) and left dipped in isopropanol until required.

### Image Analysis

Image analysis for this study was performed using Fiji [59] except traction force microscopy, monolayer stress microscopy and bayesian force inference analysis, which was performed in MATLAB (MathWorks). To measure proliferation rates, number expansion for MDCK-wildtype (WT) or -HRas^V12^ transformed (MT) live samples cultured on 4 kPa polyacrylamide (PAA) gels, starting from a similar seeding density and imaged after indicated time-points. Similarly, EdU stained fixed samples for the indicated time points, counterstained with DAPI, were imaged to quantitate the fraction of EdU^+^ (EdU^+^ DAPI^+^) to the total number of cells (EdU^+^ DAPI^+^ and EdU^-^ DAPI^+^). Cellpose [60] was used to segment and count the number of cells or the number of nuclei, respectively as well as to create segmented masks for mouse intestinal epithelium cross-sections. Paxillin puncta were counted using the Analyze particle tool in Fiji, averaged over respective cell area, and plotted. Temporal color-coded panels were prepared using the hyperstack module in Fiji.

### Statistical analysis

Statistical analyses were carried out in GraphPad Prism 9. Statistical significance was calculated by Unpaired t-test with Welch’s correction. Scatter-bar plots were displayed as mean ± s.e.m or mentioned in the respective figure legends. p-values greater than 0.05 were considered to be statistically not significant. No statistical methods were used to set the sample size. Quantification was done using data from at least three independent biological replicates. For analysis involving live-imaging experiments, data were collected from three independent experiments. All the experiments with representative images were repeated at least three times.

## Acknowledgements

We thank Simran Rawal for her assistance related to microfabrication. This work is funded by DBT/Wellcome Trust India Alliance (grant no. IA/I/17/1/503095 to T.D.). Mouse work was performed by P.G. as an ‘Infosys Fellow’ in Yasu Lab at Kyoto University, Japan (under the Infosys-TIFR Leading Edge travel grant with ref.: TFR/Efund/44/Leading Edge TG (R-9)/06/). We also acknowledge the intramural funds at Tata Institute of Fundamental Research, Hyderabad from the Department of Atomic Energy, India (under Project Identification No. RTI 4007) and generous funding by the Human Frontier Science Program (HFSP, grant no. RGP0007/2022). This work was also supported in part by the NIGMS of the National Institutes of Health (NIH) under award number R35GM15049 (D.B.), the National Science Foundation grant DMR-2046683 (D.B. and S.K.), the Center for Theoretical Biological Physics grant PHY-2019745 (D. B.) and funding from the Alfred P. Sloan Foundation (D.B).

## Author contributions

D.B. and T.D. supervised the project. P.G., S.P.P., and P.D. performed experiments. S.K. and D.B. formulated the theoretical model, performed computer simulations, and analyzed simulation results. P.G., S.P.P., and T.D. analyzed experimental results. H.K.S. contributed to the development of Gel Compression Microscopy. Mice experiments were performed by P.G. and N.T. under the supervision of Y.F. at Kyoto University. P.G., S.K., S.P.P., D.B., and T.D. wrote the manuscript. All authors edited and approved the manuscript.

## Data and materials availability

All data needed to evaluate the conclusions in the paper are present in the paper and/or the Supplementary Materials.

