## Supplementary Information for "Mechanical imbalance between normal and transformed cells drives epithelial homeostasis through cell competition"

Gupta *et al*.

This document contains:

A. **Materials and Methods**
B. **Supplementary Table 1**: Polyacrylamide hydrogel composition with different stiffness
C. **Supplementary Table 2**: Antibody and Fluorophore information
D. **Supplementary Table 3**: Plasmid information
E. **Supplementary Table 4**: pH-responsive hydrogel composition
F. **Supplementary Table 5:** Primers used for mice genotyping
G. **Theoretical Modeling**
H. **Supplementary references**
I. **Supplementary figure legends**
J. **Supplementary video legends**
K. **Supplementary Figures 1-7**

**B. Supplementary Table 1: Polyacrylamide hydrogel composition with different stiffness**

| *Elastic modulus (Young's modulus)* | *0.87 kPa* | *4 kPa* | *Source (Cat. No.)* |
|---|---|---|---|
| Milli-Q $H_2O$ (μL) | 882 | 807 | Millipore |
| 40% Acrylamide (μL) | 50 | 125 | Sigma-Aldrich (A4058) |
| 2% bis-acrylamide (μL) | 50 | 50 | Sigma-Aldrich (M1533) |
| Fluorescent beads (μL) (Optional for TFM) | 12 | 12 | Sigma-Aldrich (L3280) |
| 10% Ammonium persulfate, APS (μL) | 5.75 | 5.75 | Sigma-Aldrich (A3678) |
| N,N,N′,N′-Tetramethyl ethylenediamine, TEMED (μL) | 0.25 | 0.25 | Sigma-Aldrich (T7024) |

**C. Supplementary Table 2: Antibody and Fluorophore information**

| *S.no.* | *Antibody/Fluorophore* | *Cat.no. (Source)* | *Dilution/Working concentration* |
|---|---|---|---|

| | | | |
|---|---|---|---|
| 1. | DAPI (4′,6-diamidino-2-phenylindole) | D1306 (Invitrogen) | 1 µg $ml^{-1}$ |
| 2. | Click-iT EdU Imaging Kit | C10086 (Life Technologies) | NA |
| 3. | Anti-pMLC2(Ser19) | 3671 (CST) | 1:50 |
| 4. | Anti-Paxillin | ab32084 (Abcam) | 1:200 |
| 5. | Anti-alpha-E-catenin | C2081 (Sigma-Aldrich) | 1:2000 |
| 6. | Anti-E-cadherin | 14472 (CST) | 1:400 |
| 7. | Anti-E-cadherin | 3195 (CST) | 1:400 |
| 8. | Anti-E-cadherin (Clone ECCD-2) | M108 (Takara Bio) | 1:500 |
| 9. | Anti-E-Cadherin, DECMA-1 | MABT26 (Merck) | 5 µg $ml^{-1}$ |
| 10. | Anti-Rab5 | 3547 (CST) | 1:200 |
| 11. | Anti-Rab11 | 5589 (CST) | 1:200 |
| 12. | Anti-Rab7 | 9367T (CST) | 1:200 |
| 13. | Anti-LAMP1 | ab24170 (Abcam) | 1:200 |
| 14. | Anti-GFP | ab13970 (Abcam) | 1:500 |
| 15. | Anti-chicken IgY, AlexaFluor 488 | ab150169 (Abcam) | 1:500 |
| 16. | Anti-rabbit IgG, AlexaFluor 568 | A11036 (Invitrogen) | 1:500 |
| 17. | Anti-mouse IgG, AlexaFluor 568 | A11031 (Invitrogen) | 1:500 |
| 18. | Anti-rabbit IgG, AlexaFluor 647 | A21245 (Invitrogen) | 1:500 |
| 19. | Anti-mouse IgG, AlexaFluor 647 | A21236 (Invitrogen) | 1:500 |
| 20. | Alexa Fluor 647 Phalloidin | 8940 (CST) | 1:40 |

**D. Supplementary Table 3: Plasmid information**

| *S.no.* | *Plasmid* | *Source (Cat. No.)* |
|---|---|---|
| 1. | pcDNA4/TO/GFP-$HRas^{V12}$ | Gift from Prof. Yasuyuki Fujita |
| 2. | pcDNA4/TO/mApple-$HRas^{V12}$ | This study |
| 3. | pcDNA3.1/E-cadherin-GFP | Addgene (#28009) |

**E. Supplementary Table 4: pH-responsive hydrogel composition**

| *S.no.* | *Chemical species* | *Weight ratio* | *Source (Cat. No.)* |
|---|---|---|---|
| 1. | Acrylic acid (AA) | 4.054 | Sigma-Aldrich (147230) |
| 2. | 2-hydroxyethyl methacrylate (HEMA) | 29.286 | Sigma-Aldrich (P3932) |
| 3. | ethylene glycol dimethacrylate (EGDMA) | 0.334 | Sigma-Aldrich (335681) |
| 4. | 2,2-dimethoxy-2-phenylacetophenone (DMPA) | 1 | Sigma-Aldrich (196118) |

**F. Supplementary Table 5: Primers used for mice genotyping**

| *Mice* | *Forward* | *Reverse* |
|---|---|---|
| villin-CreER$^{T2}$ | CAAGCCTGGCTCGACGGCC | CGCGAACATCTTCAGGTTCT |
| DNMT1-CAG-loxP-STOP-loxP-HRas$^{V12}$-IRES-eGFP | CACTGTGGAATCTCGGCAGG | GCAATATGGTGGAAAATAAC |

**G. Theoretical modelling**

**1. Voronoi model:** In the Voronoi model, each cell is identified by its center $\{\boldsymbol{r_i}\}$ and its shape is determined based on the Voronoi tessellation of all cell centers. To capture the complex biomechanics of intra-cellular and inter-cellular interactions, each cell obeys an effective energy functional:

$$E_{cell} = K_A(A - A_0)^2 + K_P(P - P_0)^2 \quad (1)$$

The first elastic energy term in Voronoi-based vertex models is quadratic in cell area $A$, which results from the incompressibility of the three-dimensional cell volume and the resistance of the monolayer to height fluctuations (14, 15). The elasticity modulus $K_A$ represents the single-cell area elasticity, while $A_0$ denotes the area that a single cell prefers in its homeostatic condition. The second term, which is quadratic in cell cross-sectional perimeter $P$, is a combination of active contractility in the actomyosin subcellular cortex and effective cell-membrane tension arising from the competition between cortical tension and cell-cell adhesion (15, 16). This term gives rise to a preferred cell perimeter $P_0$, and $K_P$ represents the single-cell perimeter elasticity.

In our simulations we set $K_P = 1$ for all cells and used $A_0^{WT}$ as the unit of length. Hence, the sum total mechanical energy function (nondimensionalized) for the cell layer can be derived from the Eq. (1) as:

$$E = \sum_{i \in WT\ cells} (A_i - 1)^2 + (P_i - P_0^{WT})^2 + \sum_{i \in M\ cells} k\ (A_i - A_0^M)^2 + (P_i - P_0^M)^2 \quad (2)$$

where we have defined the ratio of cell area elasticities $k = K_A^M/K_A^{WT}$.

We can derive the mean hydrostatic cellular pressure ($\Pi$) (17) which, at the single cell level is given by:

$$\Pi_i = -2K_A^i(A_i - A_0^i) \tag{3}$$

and the mechanical tension at the "$i - j$" cell-cell junction (17, 18) as:

$$T_{ij} = [(P_i - P_0^i) + (P_j - P_0^j)] \tag{4}$$

Also, the mean interaction normal stress ($\sigma_n$) (17) at the single cell level is given by:

$$\sigma_n^i = -2\Pi_i + \frac{1}{2A_i}\sum_j T_{ij}L_{ij} = \sigma_{n,\Pi}^i + \sigma_{n,T}^i \tag{5}$$

which is a combination of stress components contributed by the internal cellular pressures ($\sigma_{n,\Pi}^i$) and the cell-edge tensions ($\sigma_{n,T}^i$) (see Supplementary Fig. 4C-D).

**2. Bulk compressibility of the tissue:** To theoretically study the effect of compressive stress on the areal densities of the two competing cell populations, we defined a relative compaction parameter $\Delta\rho$ similar to the relative compaction measurements of the experimental data reported in the main text as:

$$\Delta\rho = \frac{\rho_c - \rho_h}{\rho_h} = \frac{\rho_c}{\rho_h} - 1 = \frac{A_h}{A_c} - 1 \equiv \frac{A_0^{cell}}{\bar{A}_{cell}} - 1 \tag{6}$$

where $\rho_c$ and $A_c$ are respectively the areal densities and cell area of a cell population in the monolayer while in competition with the other cell-type, and $\rho_h$ and $A_h$ are respectively the density and area of a cell-type in their homeostatic environment. Hence, $A_h$ and $A_c$ could be equivalently written from the SPV as the preferred cell area $A_0^{cell}$ of a cell-type and the mean area $\bar{A}_{cell} = \langle A_i \rangle_{cell}$ of the cell-type in the heterogeneous monolayer. As seen in our simulations (Fig. 4C) in sync with our experimental observations (Figs. 1F, 2H), $\Delta\rho > 0$ indicates a compaction in cell area ($A_0^M > \bar{A}_M$) when the transformed cells were under compressive stress or elevated pressure, i.e., the transformed cells were struggling to reach their desired area while in competition with the wildtype, wherein the elevated average pressure in the M population ($\Pi = \langle \Pi_i \rangle_M > 0$) were driven by tuning their homeostatically preferred area ($A_0^M$).

To quantify the effect of the ratio of area elastic moduli of single cells $k = K_A^M/K_A^{WT}$ in the increased compaction experienced by the transformed population, we calculated the linear response compressibility ($C$), denoted as $C = \frac{\partial \Delta\rho}{\partial \Pi}$, which is measured in the vicinity of $\Pi = 0$ in Fig. 4C. This assumes a linear dependence of $\Delta\rho$ on $\Pi$ in the region of a small deviation away from the uniform preferred cell area condition ($A_0^M = A_0^{WT} = 1$). This essentially treats the tissue as a linearly elastic material.

**3. Active compression of the M colony and tissue-level elastic moduli:** We incorporated an active compression into the SPV by allowing the WT cells to exert an active self-propelled force on the M island. The self-propelled force in the overdamped equation of motion of the cell-centers ($\{\mathbf{r}_i\}$ ) is modulated by the length of the shared junctions between the WT and M cells as:

$$\frac{d\boldsymbol{r}_i}{dt} = \mu\boldsymbol{F}_i + \sum_{j \in neighbours\ of\ i} L_{ij}v_0\hat{\boldsymbol{n}}_{i,j} \tag{7}$$

where the cell "$i$" exerts a self-propulsion force with magnitude $v_0/\mu$ times the length of the cell-edge $L_{ij}$ that it shares with its neighbour "$j$" ($\mu$ is the mobility, or the inverse of a frictional drag) and the direction of propulsion $\hat{\boldsymbol{n}}_{i,j}$ is consistently oriented along the cell-centers (19) with:

$$v_0 \begin{cases} = 0\,, & when\ (i,j) \in same\ type\ (either\ WT \leftrightarrow WT\ or\ M \leftrightarrow M) \\ = 0\,, & when\ i \in M, j \in WT \\ \neq 0\,, & when\ i \in WT, j \in M \end{cases} \tag{8}$$

$\boldsymbol{F}_i$ is the effective mechanical interaction force experienced by the cell "$i$". In other words, only WT cells can actively push their M neighbours only, directed from WT-center towards the M-center. We chose a fixed value of $v_0 = 0.1$ for $i \in WT, j \in M$ cases, i.e., to allow the WT cells to probe an active force small in magnitude. Together these forces control the overdamped equation of motion of the cell centers.

We first allowed the heterogenous confluent layer to reach a steady state by minimising Eq. SI-2, then we studied the dynamics of the active compression by solving Eq. SI-8. At every step $(t)$ of applying active compression, we extracted the compressive areal strain as:

$$\epsilon(t) = (A_T(t) - A_T(t=0))/A_T(t=0) \tag{9}$$

by measuring the relative change in the total occupied area of the colonies $A_T(t) = \sum_i A_i(t)$ from the initial point of applying active compression. This enabled us to calculate the bulk modulus ($B$), which is the inverse of compressibility, in the linear response regime as:

$$B = \lim_{t \to 0} \frac{\partial \sigma(t)}{\partial \epsilon(t)} \tag{10}$$

where $\sigma(t) = \langle \sigma_n^i(t) \rangle$ is the average normal stress measured in the monolayer in the colonies. The differential bulk modulus between the M and WT colonies is given by $\Delta B = B_M - B_{WT}$. We employed the mean edge tension in the colonies as an approximation of the shear modulus and measured the differential tension $\Delta\tau = \tau_M - \tau_{WT}$ where:

$$\tau_{cell} = \langle P_i \rangle_{cell} - P_0^{cell} \tag{11}$$

As reported in the main text of this work, $(\Delta B - b\ \delta q_0)$ as a function of $kA_0^M$ shows collapse of data for various values of $\delta q_0$ (Fig. 5B) and $\Delta\tau/k^d$ as a function of $\delta q_0$ shows a universal scaling collapse for various values of $k$ (Fig. 5D). We found the scaling parameters to be $b = 1.7 \pm 0.2$ and $d = 0.53 \pm 0.12$ by defining error functions based on the "closeness" of the collapse (using standard deviation) and minimizing the error functions to the respective scaling parameters as variables. The errors in the measured scaling parameters are estimated by choosing a range of 5% variation in the minimized error function values.

Based on these results we proposed a phase diagram with phase boundaries determined by $\Delta B = 0$ and $\Delta\tau = 0$ to identify four regimes of mechanical cell competition, reported in the main text of this work (Fig. 5E).


### I. Supplementary figure legends

**Supplementary Figure 1. Proliferation differential characterization between wildtype and HRas$^{V12}$-transformed mutant cells. A.** Representative image panels for MDCK-wildtype (WT) or HRas$^{V12}$-transformed mutant cells, cultured for the indicated time points starting from similar seeding density in order to measure their homeostatic number expansion. Scale bar, 10 μm. **B.** Number expansion characterization for MDCK-WT or -MT cells homogenously cultured until the indicated time points. Data are mean±sem from three independent experiments. Statistical significance was calculated using Unpaired t-test with Welch's correction. **C.** Representative image panels to show EdU$^+$ cells (marks proliferating cells, as marked by EdU click-tagged with AlexaFluor-594) for MDCK-WT or -MT cells, cultured for the indicated time points starting from a similar seeding density in order to measure their homeostatic growth rate. Scale bar, 10 μm. **D.** EdU$^+$ cell fraction measurement for MDCK-WT or -MT cells homogenously cultured until the indicated time points. Data are mean±sem from three independent experiments. Statistical significance was calculated using Unpaired t-test with Welch's correction.

**Supplementary Figure 2. Experimental setup and mechanical characterization of cell competition between wildtype and HRas$^{V12}$-transformed mutant population. A.** Experimental procedure description to set-up MDCK-wildtype (WT) and HRas$^{V12}$-transformed mutant competition on a polyacrylamide (PAA) gel of defined stiffness when seeded in 40:1 (WT:transformed) number ratio. **B.** Image panels for MDCK-WT, -MT or competition between the two cell-types depicting immunostained endogenous phospho-Myosin Light Chain 2 (pMLC2) to mark contractile actomyosin. Sharp pMLC2 enrichment is seen around MDCK-MT cells when surrounded by -WT cells but not in homogenous culture conditions. Scale bar, 10 μm. **C-D.** Monolayer stress microscopy-based stress mapping in a homogenous MDCK-WT (**C**) or MDCK-MT (**D**) depicting predominant tensile stressed in such homogenous conditions for either cell-type. Scale bar, 50 μm. **E.** Time course dependent traction force microscopy and monolayer stress microscopy in a progressively competing MDCK-WT and -MT regime. This comprehensive analysis means to show how traction forces and impending stresses develop as a function of the competition time scale (incurred by the increasing time scale of HRas$^{V12}$ induction with doxycycline). Transformed colony is marked by black dotted circle. hpi- hours post induction with doxycycline. Scale bar, 50 μm.

**Supplementary Figure 3. Cell-cell junction fidelity and cell-matrix adhesion characterization in HRas$^{V12}$-transformed mutant cells.** Representative panels to show immunostained endogenous α-catenin in (**A**) MDCK-wildtype (WT) cells and its respective (**B**) intensity line-plot or (**C**) MDCK-HRas$^{V12}$-transformed mutant cells and its respective (**D**) intensity line-plot (Magenta line refers to the intensity plot line). Cells are cultured in homogenous conditions. Insets are shown (marked by red boxes) to represent one cell in each condition. Scale bar, 10 μm for all image panels and 2 μm for all insets. DAPI intensity (marked by cyan line) is plotted as reference to the cell nucleus. Pink boxes refer to the cytosol in each cell. Intensity values are normalized to the highest value within each dataset. α-catenin shows distinct junctional accumulation in MDCK-WT cell but diffused cytoplasmic signal in MDCK-MT cell. **E.** Live-image time-projected panels of fluorescently labelled plasma membranes for MDCK-WT or -MT cells. Loosely coupled membranes show long-range undulations and can be captured live which is seen as a broadly localized intensity signal in MDCK-MT cells but a collimated, sharply localized intensity signal in MDCK-WT cells and is plotted as an intensity profile in **F**, intensity values are normalized to the highest value within each dataset. Scale bar, 5 μm. **G.** Cell membrane live dynamics are visualized in MDCK-WT or -MT cells with fluorescently labelled plasma membrane and the temporal dynamics for each cell-type is shown as a colour time projection map where each time stamp is differentially coloured (in the pallet shown) and superimposed. Each time frame refers to a 5 second imaging window. **H-I.** Representative image panels depicting Paxillin based cell-matrix focal adhesions in MDCK-WT (**H**) or -MT (**I**) cells cultured in homeostatic conditions. Paxillin puncta are represented in the basal-plane of the cells while mid-plane is shown as a reference to the cells. Scale bar, 10 μm. **J**. Normalized Paxillin$^{+}$ puncta count is estimated by counting the total number of paxillin puncta in the basal-plane for both MDCK-WT and -MT using Fiji's analyze particle tool and plotted. Data are mean±sem from three independent experiments. Statistical significance was calculated using Unpaired t-test with Welch's correction.

**Supplementary Figure 4. Identifying critical factors using the SPV model. A.** The junctional tension (Eq. SI-4) at the edges shared by the transformed mutant (M) cells is significantly higher than that of their surrounding wildtype (WT) counterparts when $q_0^M\left(= P_0^M/\sqrt{A_0^M} = 3.1\right) < q_0^{WT}(= 3.65)$ . **B.** The system exhibits average normal stress in the M island (with the same spatial distribution as shown in Fig. 4B and coarse-grained over

50 ensembles of initial configurations) is lower, indicating M cells experiencing compressive stress from surrounding WT cells. **C.** Contribution of pressure in the normal stress (corresponding to the first term in the right-hand side of Eq. SI-5). **D.** Contribution of cell-edge tension in the normal stress (corresponding to the second term in the right-hand side of Eq. SI-5).

**Supplementary Figure 5. Gel Compression Microscopy (GCM). A.** Illustration to pictographically describe the setup for gel compression microscopy (GCM). Each component from the setup is separately labelled in the pictograph. **B.** Image acquired for a live GCM polyacrylamide (PAA) donut-shaped gel prepared under the micropatterned PDMS mould (referred in **Fig. 4E**). **C.** Creep analysis is plotted where compressive strain (as fraction to the pre-compression perpendicular shift of the interface) incurred during GCM is plotted as a function of the corresponding time of compression in minutes. The total duration for compression is 240 minutes where at about 90 minutes of compression, there is a sharp non-linearity in compression for the MDCK-WT population but not MDCK-HRas$^{V12}$-transformed mutant population or PAA gel alone condition. Data are mean±sd from three independent experiments. **D**. Relative compressibility quantitation derived mean±sem values depicted in tabular form for experiments with 4 kPa polyacrylamide substrate for all conditions. **E.** Relative compressibility quantitation derived mean±sem values depicted in tabular form for experiments with 0.87 kPa polyacrylamide substrate for all conditions. Data are mean±sem from three independent experiments. Statistical significance was calculated using Unpaired t-test with Welch's correction.

**Supplementary Figure 6. Characterization of E-cadherin endocytic trafficking in HRas$^{V12}$-transformed mutant cells.** Representative image panels for MDCK-wildtype (WT) or MDCK-HRas$^{V12}$-transformed mutant cells stained for endogenous E-cadherin and counterstained for **A,** Rab5 for early endosomes, **B,** Rab11 for recycling endosomes, **C,** Rab7 for late endosomes, and **D,** LAMP1 for lysosomes. Insets are shown over respective images (marked by white boxes). Cyan arrowheads show E-cadherin$^{+}$ vesicles whereas white arrowheads show respective endocytic-marker$^{+}$ compartment. In either MDCK-WT or -MT case, E-cadherin did not show preferential co-localization with any of the endocytic compartments. Scale bar, 5 μm for all image panels and 2 μm for all insets. A pictograph depicting the general E-cadherin endosomal trafficking route in living cells is shown on the *right*.

**Supplementary Figure 7. Gel Compression Microscopy (GCM) based collective compressibility measurements for E-cadherin rescue HRas$^{V12}$-transformed mutant population. A.** Representative image panels for 4 kPa PAA gel control, MDCK-wildtype (WT) cell monolayer or MDCK-HRas$^{V12}$-transformed mutant cells or MDCK-HRas$^{V12}$ transformed cells overexpressing E-cadherin-GFP, cultured on 4 kPa PAA gels for GCM. Pre-compression ROIs are shown to depict PAA gel alone or cells-PAA gel interface (marked by yellow dotted lines) as well as their respective post-compression ROIs representing the respective interfaces after 90 mins of continuous compression (marked by white dotted lines), Scale bar, 50 μm. **B**. Relative compressibility quantitation derived mean±sem values depicted in tabular form for experiments with 4 kPa polyacrylamide substrate for all conditions. Data are mean±sem from three independent experiments. Statistical significance was calculated using Unpaired t-test with Welch's correction.

### J. Supplementary video legends

**Supplementary video 1:** GFP-HRas$^{V12}$ (*top left panel*), cell dynamics (DIC, *top right panel*), traction dynamics (bottom left panel) and monolayer stress dynamics (*bottom right panel*) in a competing heterogenous monolayer of MDCK-wildtype (WT) and MDCK-HRas$^{V12}$ (MT) cells (40:1 ; WT:MT). The movie is shown at five frames per second. Time interval, 1 hour. Scale bar, 50 µm.

**Supplementary video 2:** Gel Compression Microscopy with a 4 kPa polyacrylamide hydrogel (labelled with 0.5µm orange beads) interfacing a pH-responsive hydrogel. Images are acquired in differential interference contrast (DIC). The movie is shown at five frames per second. Time interval, 1 minute. Scale bar, 50 µm.

**Supplementary video 3:** Gel Compression Microscopy with a MDCK-wildtype monolayer of cells cultured atop a 4 kPa polyacrylamide hydrogel and interfacing a pH-responsive hydrogel. Images are acquired in differential interference contrast (DIC). The movie is shown at five frames per second. Time interval, 1 minute. Scale bar, 50 µm.

**Supplementary video 4:** Gel Compression Microscopy with a MDCK-HRas$^{V12}$ monolayer of cells cultured atop a 4 kPa polyacrylamide hydrogel and interfacing a pH-responsive hydrogel. Images are acquired in differential interference contrast (DIC). The movie is shown at five frames per second. Time interval, 1 minute. Scale bar, 50 µm.

**Supplementary video 5:** Gel Compression Microscopy with a 0.87 kPa polyacrylamide hydrogel (labelled with 0.5µm orange beads) interfacing a pH-responsive hydrogel. Images are acquired in differential interference contrast (DIC). The movie is shown at five frames per second. Time interval, 1 minute. Scale bar, 50 µm.

**Supplementary video 6:** Gel Compression Microscopy with a MDCK-wildtype monolayer of cells cultured atop a 0.87 kPa polyacrylamide hydrogel and interfacing a pH-responsive hydrogel. Images are acquired in differential interference contrast (DIC). The movie is shown at five frames per second. Time interval, 1 minute. Scale bar, 50 µm.

**Supplementary video 7:** Gel Compression Microscopy with a MDCK-HRas$^{V12}$ monolayer of cells cultured atop a 0.87 kPa polyacrylamide hydrogel and interfacing a pH-responsive hydrogel. Images are acquired in differential interference contrast (DIC). The movie is shown at five frames per second. Time interval, 1 minute. Scale bar, 50 µm.

**Supplementary video 8:** E-cadherin-GFP (*yellow*) dynamics in either a MDCK-wildtype or a MDCK--HRas$^{V12}$ transformed cell. The movie is shown at five frames per second. Time interval, 5 seconds. Scale bar, 2 µm.

**Supplementary video 9:** Gel Compression Microscopy with a MDCK-HRas$^{V12}$ monolayer of cells constitutively overexpressing E-cadherin-GFP, cultured atop a 4 kPa polyacrylamide hydrogel and interfacing a pH-responsive hydrogel. Images are acquired in differential interference contrast (DIC). The movie is shown at five frames per second. Time interval, 1 minute. Scale bar, 50 µm.

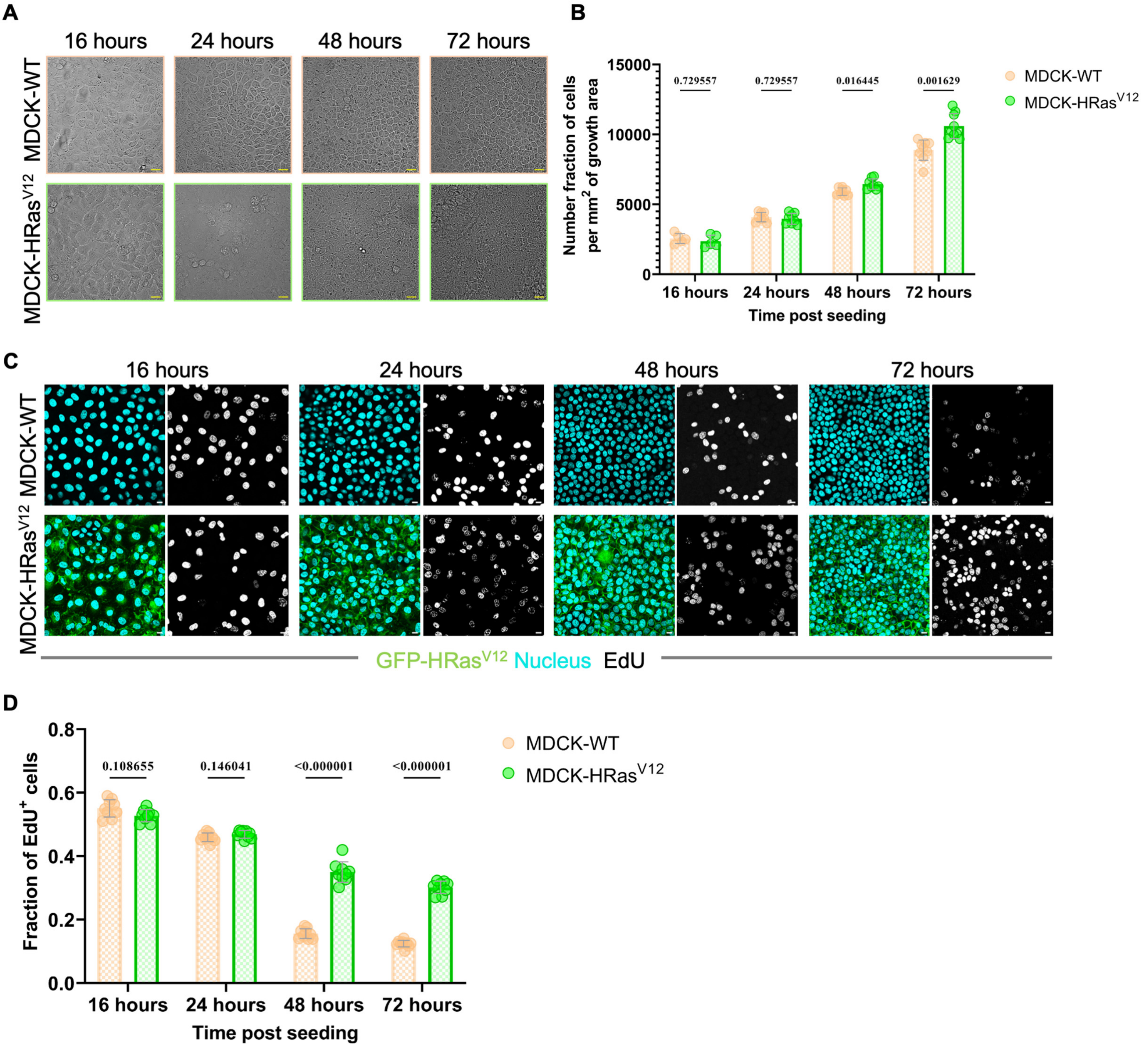


**Supplementary Figure 1**

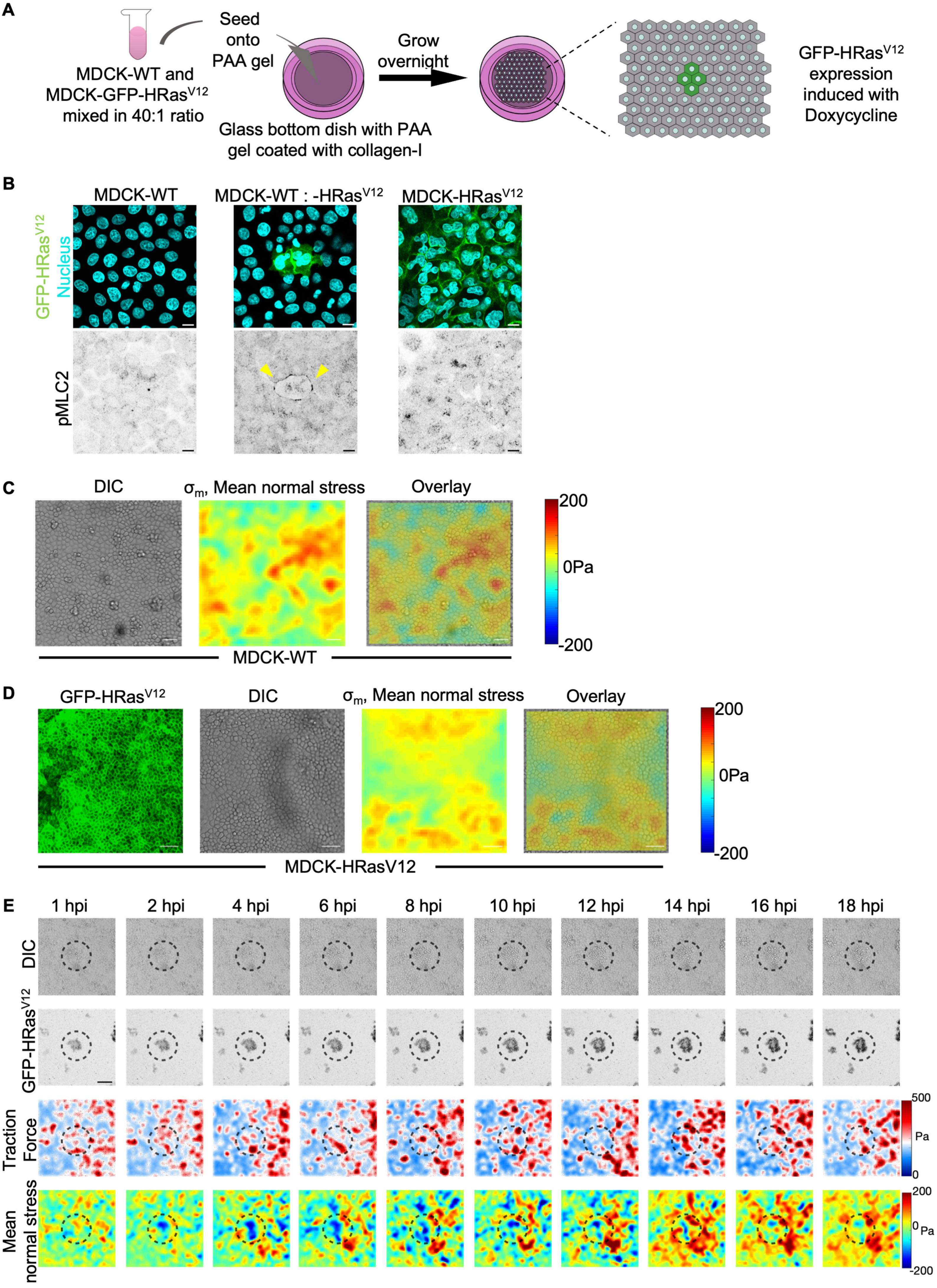


**Supplementary Figure 2**

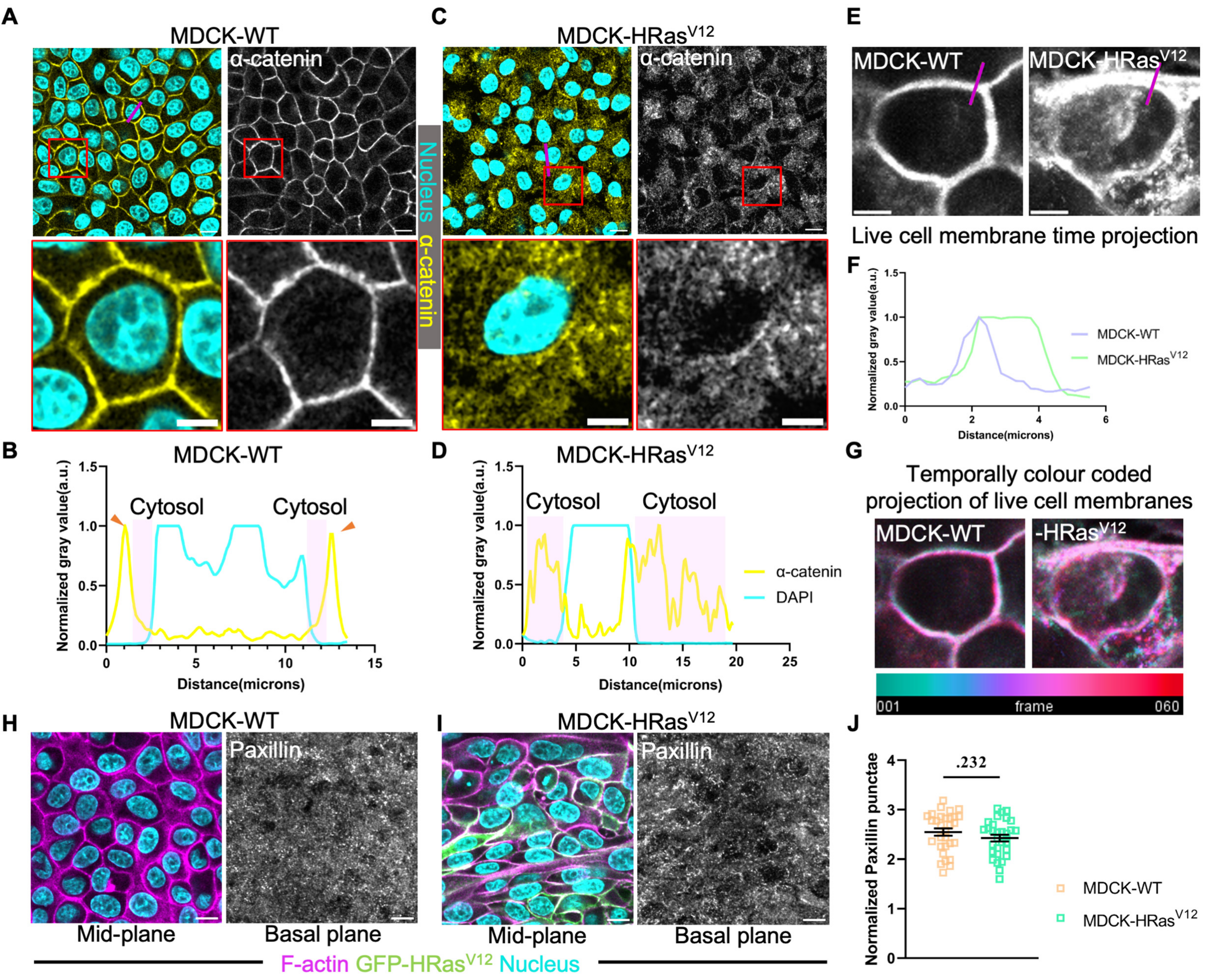


**Supplementary Figure 3**

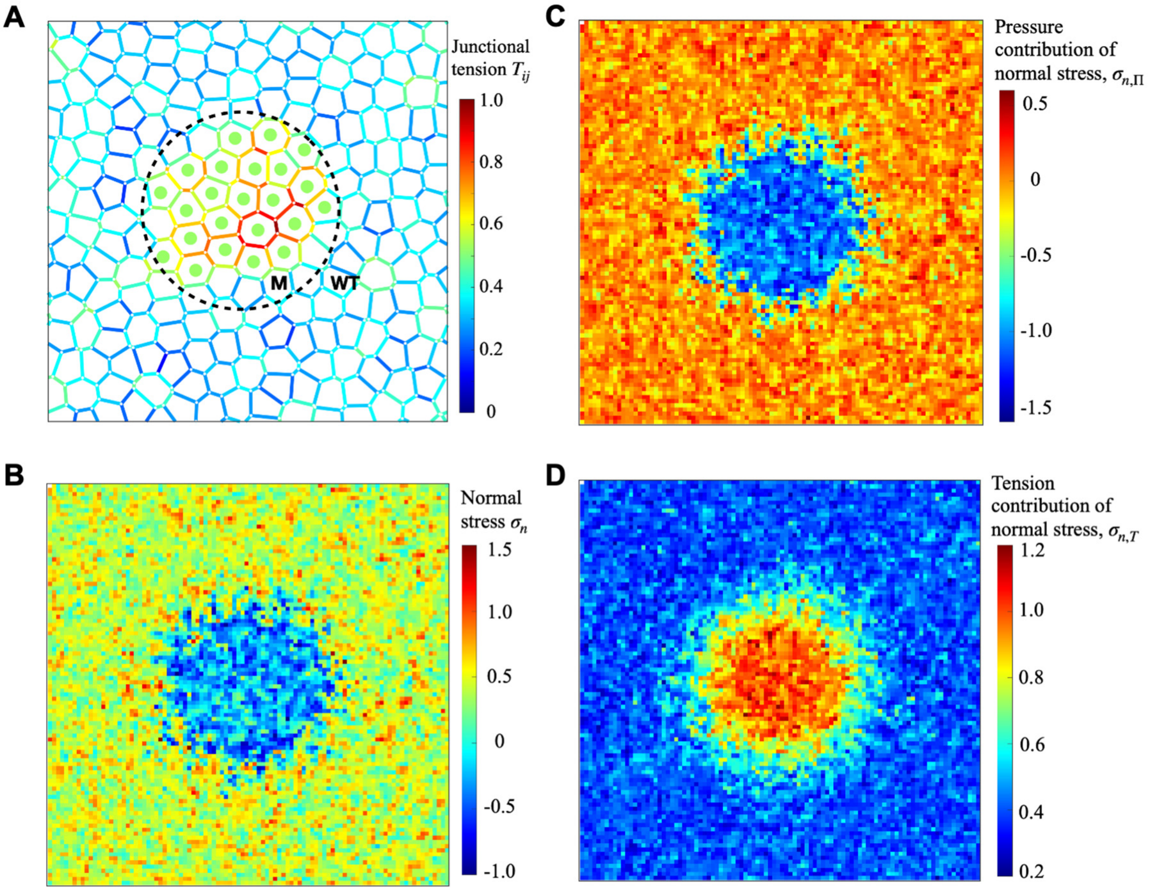


**Supplementary Figure 4**

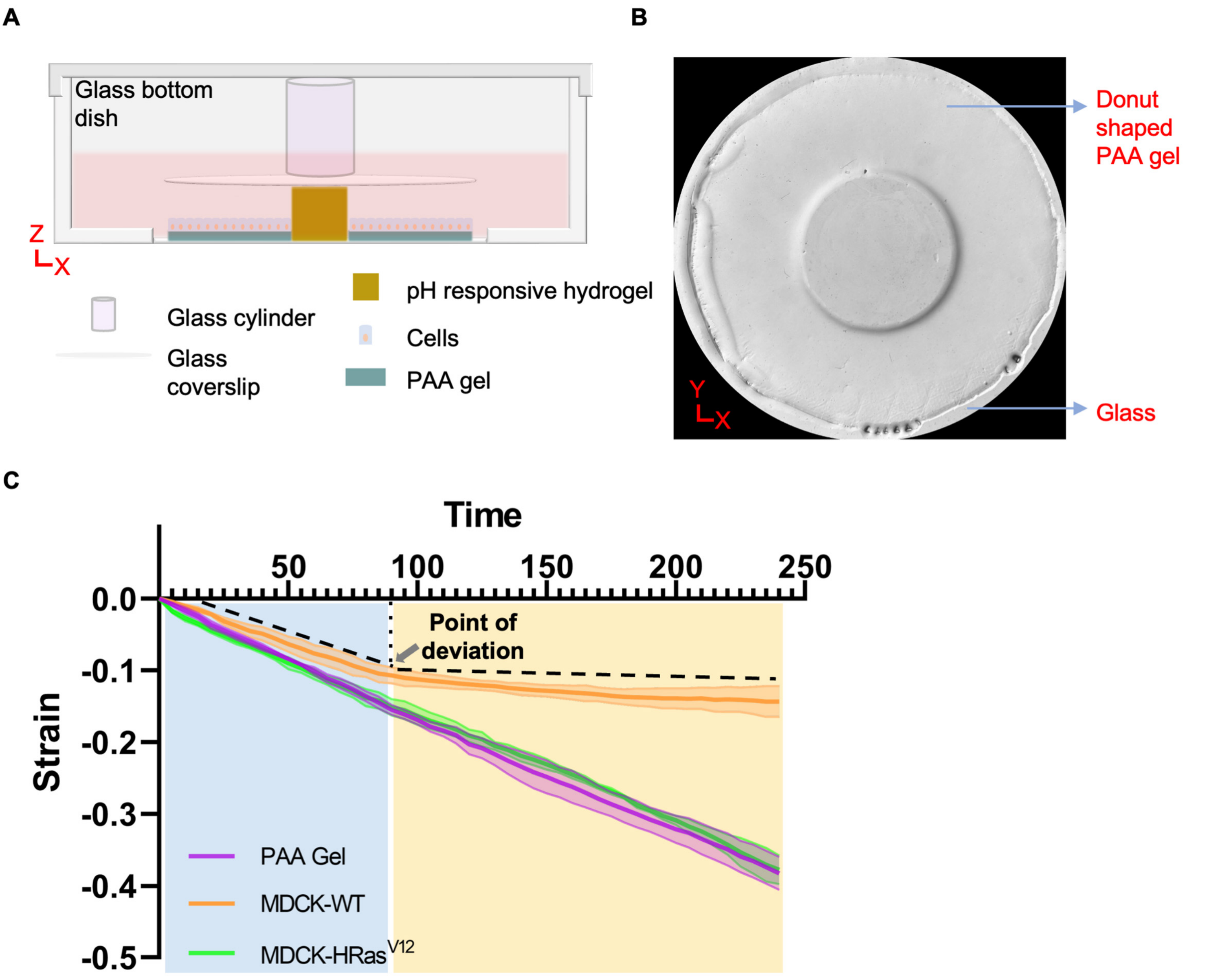


D

| Relative Compressibility (Mean ± SEM) | | |
|---|---|---|
| 4 kPa PAA Gel | MDCK-WT | MDCK-HRas$^{V12}$ |
| 1.00±0.17 | 0.49±0.07 | 0.99±0.08 |

E

| Relative Compressibility (Mean ± SEM) | | |
|---|---|---|
| 0.87 kPa PAA Gel | MDCK-WT | MDCK-HRas$^{V12}$ |
| 1.00±0.13 | 0.31±0.03 | 0.63±0.05 |

**Supplementary Figure 5**

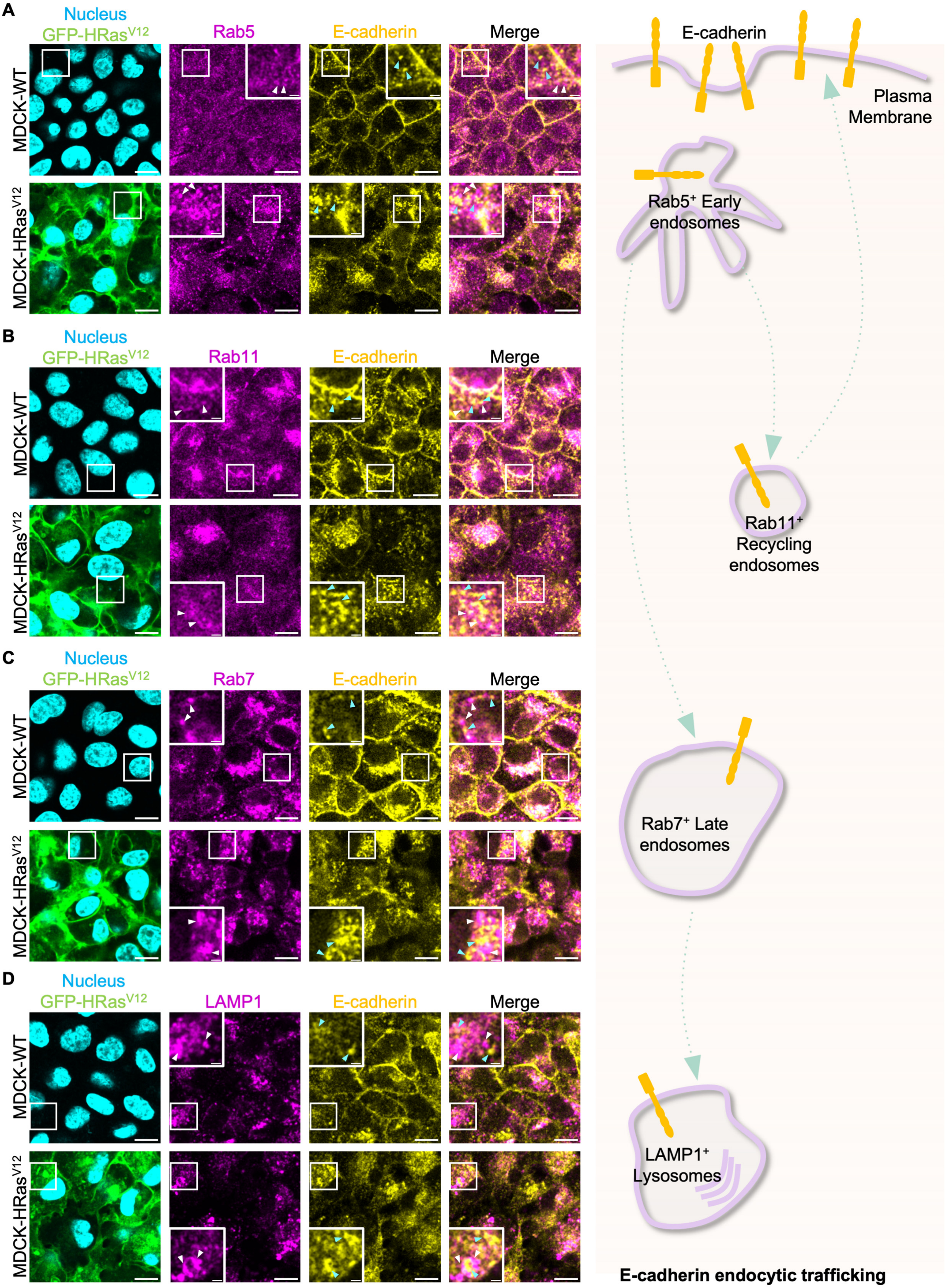


**Supplementary Figure 6**

**A**

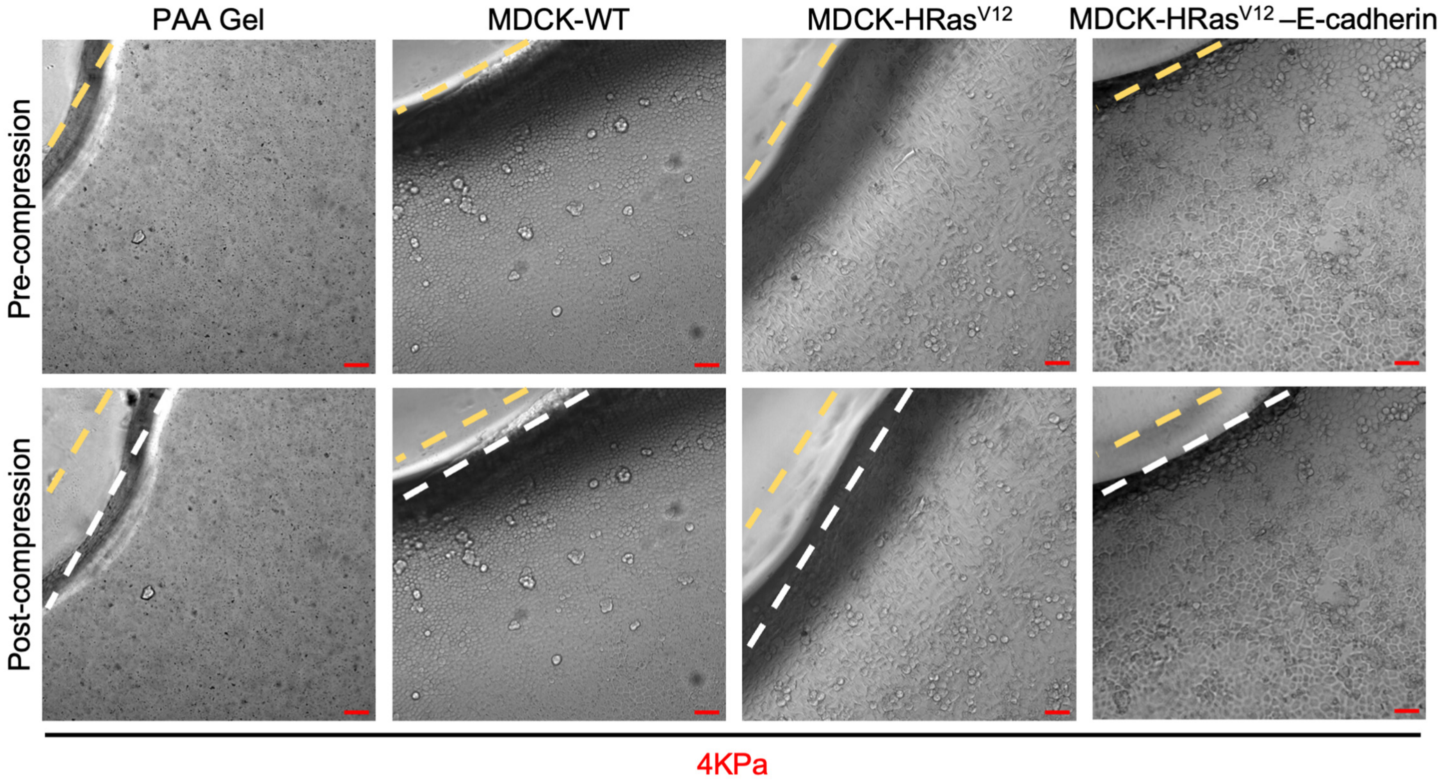


**B**

| Relative Compressibility (Mean ± SEM) | | | |
|---|---|---|---|
| 4 kPa PAA Gel | MDCK-WT | MDCK-HRas$^{V12}$ | MDCK-HRas$^{V12}$ – E-cadherin |
| 1.00±0.14 | 0.53±0.02 | 1.01±0.10 | 0.53±0.02 |

**Supplementary Figure 7**
